# A mouse model of human mitofusin 2-related lipodystrophy exhibits adipose-specific mitochondrial stress and reduced leptin secretion

**DOI:** 10.1101/2022.09.20.508662

**Authors:** JP Mann, X Duan, A Alvarez-Guaita, A Haider, I Luijten, M Page, S Patel, F Scurria, M Protasoni, LC Tábara, S Virtue, S O’Rahilly, M Armstrong, J Prudent, RK Semple, DB Savage

**Affiliations:** University of Cambridge Metabolic Research Laboratories, Wellcome Trust-MRC Institute of Metabolic Science, Cambridge, CB2 0QQ, UK; Centre for Cardiovascular Science, University of Edinburgh, Edinburgh, UK; UCB Biopharma UK, Bath Road, Slough, UK; Medical Research Council Mitochondrial Biology Unit, University of Cambridge, Cambridge, UK; UCB Pharma, Chemin du Foriest 1, 1420 Braine l’Alleud, Belgium; MRC Human Genetics Unit, Institute of Genetics and Cancer, University of Edinburgh, Edinburgh, UK

**Keywords:** Mitochondria, mitofusin, adipose tissue, integrated stress response, leptin

## Abstract

Mitochondrial dysfunction has been reported in obesity and insulin resistance, but primary genetic mitochondrial dysfunction is generally not associated with these, arguing against a straightforward causal relationship. A rare exception, recently identified in humans, is a syndrome of lower body adipose loss, leptin-deficient severe upper body adipose overgrowth, and insulin resistance caused by the p.Arg707Trp mutation in *MFN2*, encoding mitofusin-2. How this selective perturbation of mitochondrial function leads to tissue- and adipose depot-specific growth abnormalities and systemic biochemical perturbation is unknown. To address this, *Mfn2^R707W/R707W^* knock-in mice were generated and phenotyped on chow and high fat diets. Electron microscopy revealed adipose-specific mitochondrial morphological abnormalities. Oxidative phosphorylation by isolated mitochondria was unperturbed, but the cellular integrated stress response was activated in adipose tissue. Fat mass and distribution, body weight, and systemic glucose and lipid metabolism were unchanged, however serum leptin and adiponectin concentrations, and their secretion from adipose explants were reduced. Pharmacological induction of the integrated stress response in wild-type adipocytes also reduced secretion of leptin and adiponectin, suggesting an explanation for the *in vivo* findings. These data suggest that the p.Arg707Trp MFN2 mutation perturbs mitochondrial morphology and activates the integrated stress response selectively in adipose tissue. In mice, this does not disrupt most adipocyte functions or systemic metabolism, whereas in humans it is associated with pathological adipose remodelling and metabolic disease. In both species, disproportionate effects on leptin secretion may relate to cell autonomous induction of the integrated stress response.

## Introduction

Mitochondrial dysfunction has been implicated in the pathogenesis of a wide range of congenital and acquired conditions^1–3^. However, despite being central to cellular energy homeostasis, there has been little mechanistic evidence of a causal role for deranged mitochondrial function in human adiposity. Instead, most patients with inherited mitochondrial disorders have a neurological phenotype, though multisystem involvement is common^1, 4^. This is true of disorders caused by many different mutations affecting mitochondrial proteins, whether encoded in mitochondrial or nuclear DNA. The mechanisms underlying such tissue-selective disease manifestations even in the face of constitutional mutations are unclear, but differing tissue requirements for oxidative phosphorylation, and interactions between the nuclear and mitochondrial genome may play a role^5–7^.

We^8^ and others^9–13^ recently identified biallelic R707W mutations in the nuclear *MFN2* gene in patients with a remarkable adipose phenotype characterised by extreme upper body adiposity (lipomatosis) and lower body lipodystrophy. Affected patients also showed non-alcoholic fatty liver disease, dyslipidaemia and insulin resistant type 2 diabetes, likely secondary to the changes in adipose tissue. *MFN2* encodes mitofusin 2, a mitochondrial outer membrane protein that plays a key role in mitochiondrial fusion and tethering to other organelles. Like patients with heterozygous complete loss-of-function mutations in *MFN2,* patients with the R707W mutation also often exhibit axonal peripheral neuropathy known as Charcot-Marie Tooth type 2A (CMT2A)^14–16^, but the adipose and metabolic phenotype has uniquely been associated with the R707W allele to date. Evidence suggesting that the MFN2 R707W mutation does disrupt mitochondrial function in humans includes elevated serum lactate in affected patients, abnormal mitochondrial morphology seen on transmission electron microscopy of affected adipose tissue^8^, and strong transcriptomic signatures of mitochondrial dysfunction and activation of the integrated stress response^4^ in the same tissue.

Despite normal or raised whole body adipose mass, and the relatively normal histological appearance of lipomatous upper body fat, plasma leptin concentrations were very low in the patients reported by Rocha *et al*.^8^. This observation was supported by Sawyer *et al.* who reported one patient with undetectable serum leptin (<0.6 ng/mL)^9^ and Capel *et al.* who described five patients with serum leptin concentrations <1.6ng/mL despite BMIs within the normal range^10^. Leptin is a critical endocrine signal of adipose stores and yet what determines the rate of adipocyte leptin secretion remains poorly understood^17^. This surprising observation thus offered a rare opportunity to address this important issue. Circulating leptin concentration correlates with fat mass, and is usually higher in women than men, with some adipose depots reported to release more than others^18^. Leptin secretion is increased by insulin stimulation^19^ but this effect is modest compared to the overall circulating levels. Adipose depots that secrete higher leptin have increased *LEP* mRNA and, at least *in vitro*, *LEP* mRNA increases in response to insulin stimulation^20^.

Most neuropathy-associated *MFN2* mutations are located within the protein’s GTPase domain^21–24^, but to date, all patients with *MFN2*-associated multiple symmetric lipomatosis (MSL) have had at least one R707W allele. Most cases have been homozygous for the R707W mutation, with others compound heterozygous for R707W and a second, functionally null mutation^8–13^. Arginine 707 lies in the highly conserved heptad-repeat (HR)2 region of MFN2 but consensus on the precise orientation and/or function of this domain has not yet been established^25^. The dominant model holds that the HR2 domains of Mitofusin 1 (MFN1) and/or MFN2 face the cytosol and interact in *trans,* forming mitofusin homodimers or heterodimers required for mitochondrial fusion and tethering to other organelles^26, 27^. The R707W mutation may disrupt this binding and subsequent MFN oligomerisation and mitochondrial fusion/tethering.

Global knock-out of the core mitochondrial fusion-fission machinery proteins *Mfn1*, *Mfn2*, *Opa1*, and *Drp1* in mice confers embryonic lethality in all cases^28–30^. Two homozygous loss-of-function *Mfn2* knock-ins - H361Y and R94W - have also been reported. H361Y was also embryonically lethal due to complete loss of detectable Mfn2 protein^31^, while homozygous R94W mice died at post-natal day 1^32^. Mfn2^R94W^ is expressed but is a GTPase defective mutant that increases mitochondrial fragmentation and prevents formation of Mfn2 homodimers^33^.

Multiple tissue-specific knock-outs of *Mfn2* (and/or *Mfn1*) have been studied including three in adipose tissue^34–37^. Adipose-specific *Adipoq*-Cre *Mfn2* knock-out increased fat mass, attributed to reduced energy expenditure, whether it occurred in embryonic life^35^ or was induced by tamoxifen in adult mice^37^, with ultrastructural evidence of more rounded mitochondria with fewer lipid droplet contacts^35^. Both Adipoq-Cre and *Ucp1*-Cre (brown adipose-specific) *Mfn2* knock-out also caused cold intolerance with “whitened” brown adipose tissue^36^. Plasma leptin concentrations were not reported in Adipoq-Cre or *Ucp1*-Cre *Mfn2* knock-outs, but leptin concentration was higher and adiponectin concentration lower in the tamoxifen-inducible adult Adipoq-Cre *Mfn2* knockout model^37^.

These studies suggest that *Mfn2* has a non redundant role in adipose tissues, but findings to date are not readily reconcilable with the phenotype observed in humans with the MFN2^R707W^ mutation. We thus generated and characterised mice homozygous for Mfn2^R707W^ to determine the extent of tissue-specific manifestations of mitochondrial dysfunction and to interrogate the effect of this mutant on adipose leptin secretion.

## Results

### Generation of Mfn2^R707W/R707W^ mice

*Mfn2^R707W/R707W^* mice were generated by CRISPR-Cas9 genomic engineering using an ssODN (single-stranded oligo donor) template recoding Arginine to Tryptophan in codon 707 (**Figure 1A**). A single round of targeting yielded one founder (F0) mouse (**Figure 1B**) which was used to expand a colony on a C57/BL6J background. An additional silent mutation introducing an EcoRV restriction site was introduced to facilitate genotyping (**Figure 1C, D**). We first compared expression of Mfn2 and its paralogue, Mfn1, in knock-in (KI) mice and wild-type (WT) littermates in order to determine if the R707W change perturbed expression of the mutant Mfn2 protein and/or resulted in a compensatory change in Mfn1. We observed no consistent differences in Mfn1 or Mfn2 expression, relative to WT, in white adipose tissue (WAT), liver, heart, or skeletal muscle in both chow and high fat diet (HFD) fed mice (**Figure 1E, Figure 1-Figure supplement 1 and 2**). However, in brown adipose tissue (BAT) expression of Mfn1 was lower in KI than in WT mice fed a chow diet (**Figure 1-Figure supplement 1D**) and, in HF fed mice, both Mfn1 and Mfn2 expression was lower in KI mice (**Figure 1E, Figure 1-Figure supplement 1D 2D**). We interpret these data as suggesting that the R707W missense mutation does not directly reduce expression of Mfn2, nor does it result in a compensatory change in Mfn1 in most tissues. However, in BAT the data suggests that the cellular perturbation induced in brown adipocytes is associated with a very modest reduction in expression of Mfn1 and 2.

**Figure 1.**
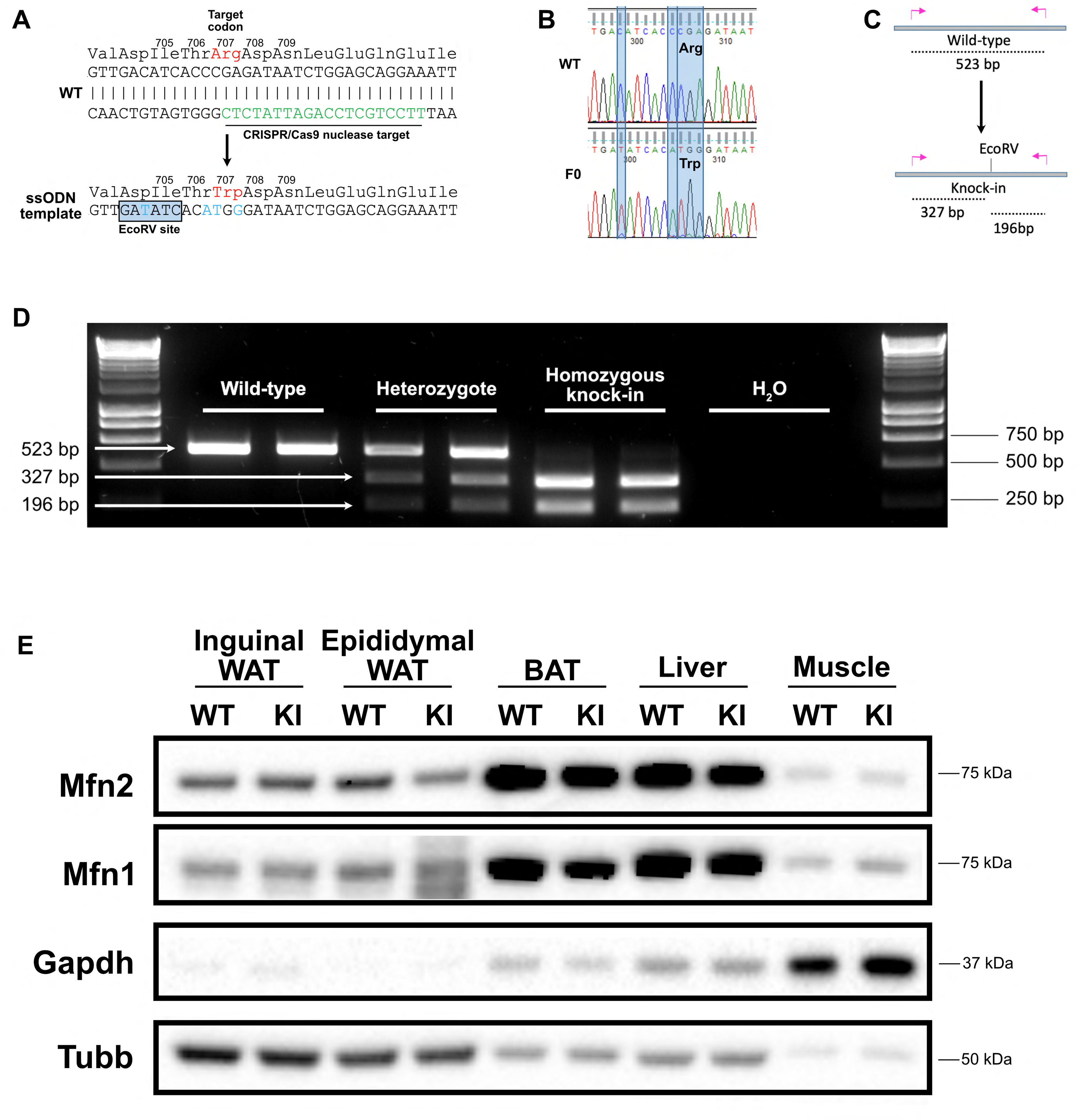
Generation of a Mfn2^R707W^ knock-in mouse. (A) Wild-type (WT) nucleotide and amino acid sequence around the Arg 707 codon. The CRISPR/Cas9 nuclease target is indicated in green. Below is part of the ssODN template with mutated nucleotides in blue, including the upstream silent mutation (at codon 704-705) to generate an EcoRV restriction site. (B) Sanger sequencing confirmation of the knock-in (KI) with restriction site in a founder (F0). (C) Illustration of the genotyping strategy: mutant alleles will digest into 327 bp and 196 bp fragments in response to EcoRV digestion. (D) Ear biopsies were digested using chelix and *Mfn2* amplified by PCR, then digested using EcoRV. Representative SYBR Safe DNA gel demonstrating genotyping for two WT, heterozygous, and homozygous KI mice. Image is representative of other genotyping gels. (E) Western blot from inguinal and epididymal white adipose tissue (WAT), brown adipose tissue (BAT), liver and skeletal muscle for expression of Mfn1 and Mfn2. Tissues are from mice fed a 45% kcal high fat diet (HFD) for 6 months. Due to variability across tissues, both *Gapdh* and Beta-tubulin (*Tubb*) are given as loading controls. The image is representative of at least three biological replicates. **Figure 1-source data.** Raw and annotated Western blots for each protein from Figure 1E.

### Mfn2^R707W/R707W^ mice show adipose-selective alterations of mitochondrial structure and function

Next, we used transmission electron microscopy (TEM) to examine mitochondrial ultrastructure **(Figure 2A)**. In BAT, mitochondria from KI mice had a tendency to exhibit a decreased mitochondrial perimeter compared to WT mice (**Figure 2B**), but significantly reduced elongation, assessed by the mitochondrial length / width aspect ratio of cross sections (**Figure 2C**), indicating that Mfn2^R707W^ leads to mitochondrial fragmentation in BAT. Double membrane-bound structures representing autophagosomes consistent with mitophagy were observed in lipomatous adipose tissue from human patients with the MFN2^R707W^ mutation^8^, but these were not identified in the murine tissues examined. Mitofusins may mediate contact between mitochondria and lipid droplets^35^, and the extent of these contacts was reduced in BAT from KI animals (**Figure 2D**). There was no difference in the number of cristae per mitochondrion (**Figure 2E**). In both inguinal (**Figure 2-figure supplement 1A-C**) and epididymal WAT (**Figure 2-figure supplement 1D-F**), similar mitochondrial fragmentation were observed. In contrast, no change in mitochondrial morphology was seen in samples from the heart, skeletal muscle or livers of KI mice (**Figure 2-figure supplement 2**).

**Figure 2.**
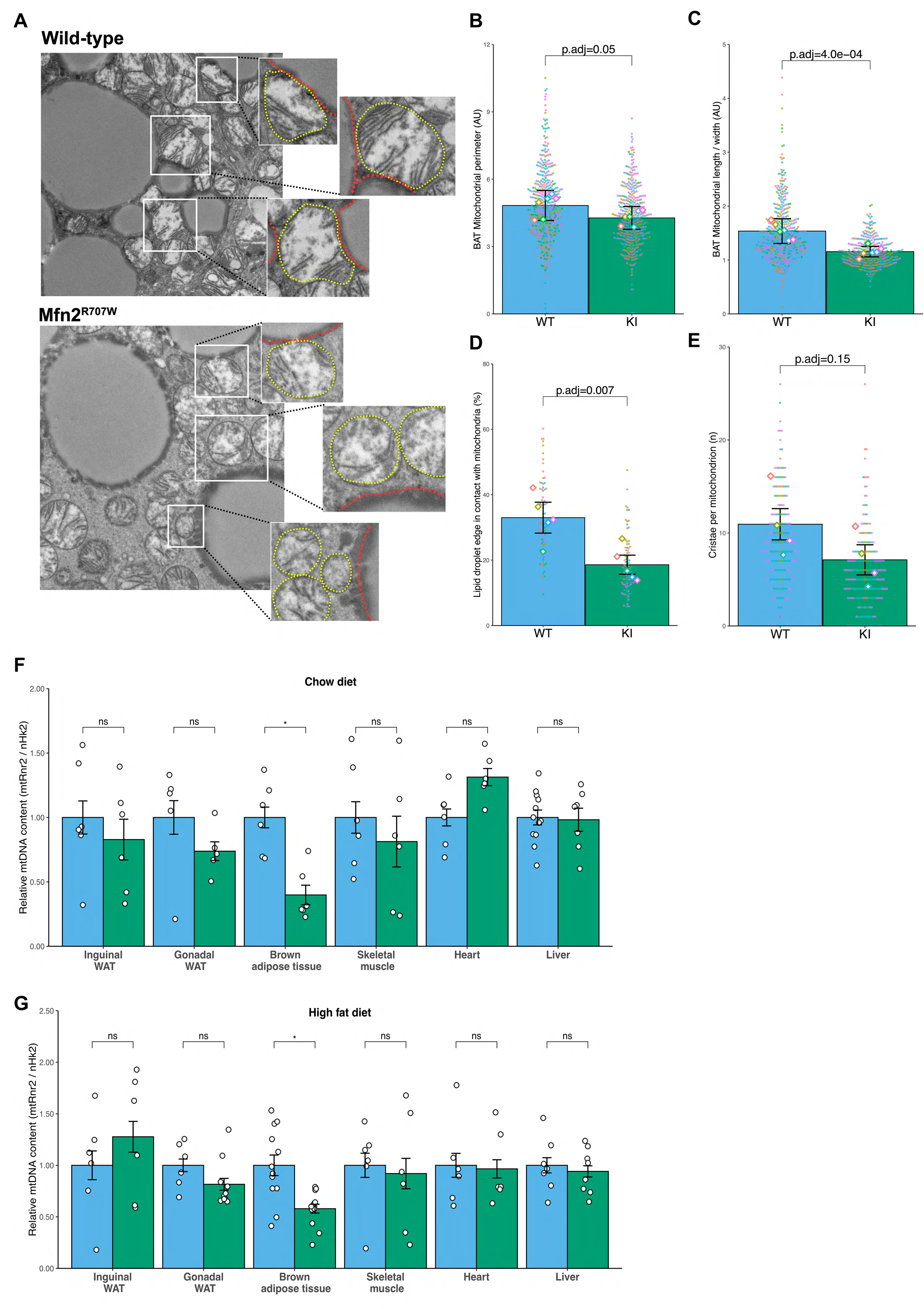
Effect of Mfn2^R707W^ on mitochondrial structure and function. (A) Representative transmission electron microscopy (TEM) images of brown adipose tissue (BAT) with zoomed-in images of mitochondria (highlighted in yellow) bordering lipid droplets (outlined in red). (B) Quantification of mitochondrial perimeter from TEM on BAT. Each dot represents data from an individual mitochondrial cross section with each diamond showing the separate 6 biological replicates. p.adj gives the false-discovery rate (FDR) adjusted p-value from across all TEM analyses. (C) Quantification of mitochondrial aspect ratio (length/width) from TEM of BAT. (D) Quantification of mitochondrial-lipid droplet contact from TEM, expressed as proportion (%) of lipid droplet in contact with mitochondrial membrane on BAT. (E) Number of cristae per mitochondrion from TEM of BAT. (F) Mitochondrial DNA content in tissues from mice fed chow or HFD (G) for 6 months. Each data point represents a separate animal. * = FDR-adjusted p-value <.05. WT in blue, homozygous Mfn2^R707W^ in green.

Perturbed mitochondrial dynamics have been associated with decreased mitochondrial DNA (mtDNA) content, as replication of mtDNA relies on balanced fusion and fission events^38^. In Mfn2^R707W/R707W^ mice, mtDNA was only reduced in BAT, but not in all other tissues analysed (WAT, heart, skeletal muscle, or liver (**Figure 2F-G**)). Immunoblotting of electron transport chain components showed reductions in Ndufb8 (complex I) and Mtoc1 (complex IV) in BAT and inguinal WAT (**Figure 2-Figure supplement 3A-B)**. These changes were more striking in epididymal WAT (**Figure 2-Figure supplement 3C**) but were not observed in liver or heart (**Figure 2-Figure supplement 3D-E)**. Uqcrc2 (complex III) expression was lower in inguinal WAT from KI animals (**Figure 2-Figure supplement 3A)**, though this was not replicated in other tissues.

To determine if these changes altered mitochondrial oxidative phosphorylation, we assessed oxidative capacity in freshly isolated mitochondria from BAT and liver by high resolution respirometry using Oroboros Oxygraphy. No significant differences were detected between WT and KI mitochondria (**Figure 2-figure supplement 4A-B**). We further assessed mitochondrial function in BAT *in vivo* by challenging mice with noradrenaline in cold (10 degrees) or thermoneutral (30 degrees) conditions to determine maximum thermogenic capacity. Again, despite a trend towards reduced thermogenic capacity in KI animals, the difference was not significant, and both groups manifested the expected increase in energy expenditure at 10°C (**Figure 2-figure supplement 4C-G**).

### Body composition and metabolic phenotype of Mfn2^R707W/R707W^ mice

We next assessed whether *Mfn2^R707W/R707W^* mice phenocopy the severe abnormal adipose topography and metabolic abnormalities of patients harbouring the same mutations. Male mice fed with either chow or HF diet for up to 6 months were assessed. Whole body mass and composition, and masses of individual adipose depots and other organs were similar in KI and WT mice throughout the study period (**Figure 3A-C & Figure 3-figure supplement 1A-D**). Moreover, no difference in hepatic steatosis or lipid droplet size was detected histologically in BAT or WAT (**Figure 3D-E & Figure 3-figure supplement 1F-G**). In keeping with the normal body composition, fasting serum glucose, insulin, triglycerides, cholesterol, lactate, and liver transaminase concentrations showed no difference between WT and KI mice (**Figure 3F-I & Figure 3-figure supplement 1E, H**). Dynamic testing of glucose and insulin tolerance was also similar between genotypes (**Figure 3G-K**).

**Figure 3.**
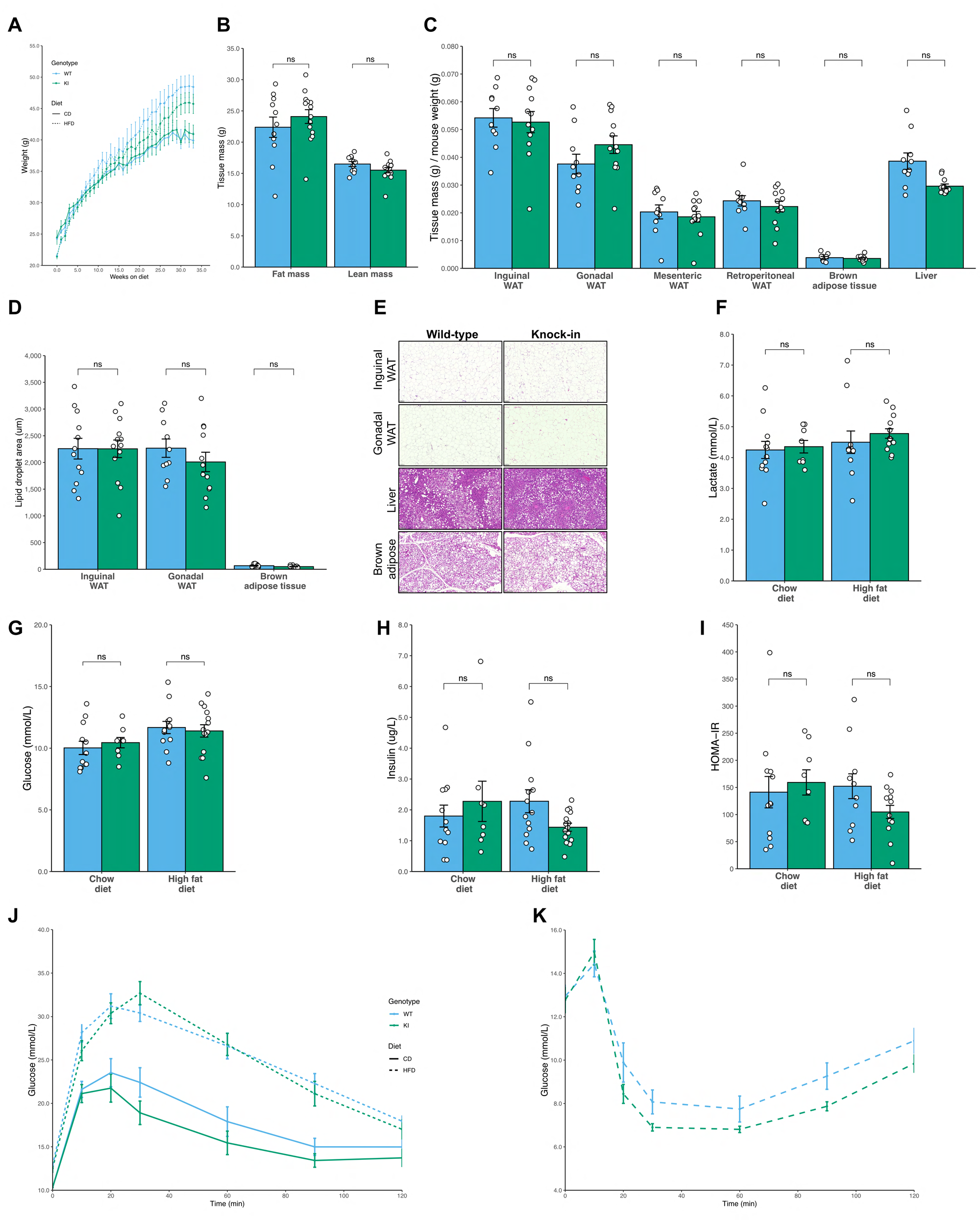
No difference in fat mass or glucose homeostasis in Mfn2^R707W^ mice on chow or high fat diet. Seven-week-old mice were fed chow (n=8-12) or 45% kcal HFD (n=13-14) for 6 months. (A) Absolute body mass for mice fed chow (solid line) and HFD (dashed line) over 6 months. (B) Time-domain nuclear magnetic resonance (TD-NMR) measurement of fat and lean mass of mice fed HFD. (C) Weights of tissues, including four white adipose tissue (WAT) depots, from mice fed HFD. (D) Quantification of lipid droplet area from histological specimens of adipose tissue from mice fed HFD for 6 months. (E) Representative histological images from WAT, liver, and brown adipose tissue. Analyses of mouse plasma lactate (F), plasma glucose (G), and insulin (H), homeostatic model of assessment of insulin resistance (HOMA-IR, I) after a 6 hr fast. Change in plasma glucose during intraperitoneal glucose tolerance test (J) and intraperitoneal insulin tolerance test (K). ns, p>.05 on un-paired T-test, adjusted for multiple comparisons. WT in blue, homozygous Mfn2^R707W^ in green. Each data point represents an individual animal.

### Adipose-tissue specific activation of the integrated stress response in Mfn2^R707W^ mice

Mitochondrial dysfunction is sensed by cells, and triggers a series of adaptive responses to maintain mitonuclear balance and cellular homeostasis^39^. Precise sensing and transducing mechanisms vary among different forms of mitochondrial perturbation^40^, and show some redundancy^41^. Although details of the integration of these mechanisms in different tissue and cellular contexts are not fully elucidated, it is clear that the transcription factor Atf4 plays a crucial role^42^. Atf4 is translationally upregulated following phosphorylation of eIF2α, which is a point of convergence of several cellular stress sensing pathways. The canonical eIF2α kinase HRI is most closely implicated in linking mitochondrial dysfunction to eIF2α phosphorylation^43, 44^, but mTORC1 appears to play a role in Atf4 upregulation independently of eIF2α phosphorylation^45^. The response to sustained mitochondrial dysfunction shares commonality with the response to many other cellular stressors, and is best characterised as a cellular integrated stress response (ISR)^46^.

Strong transcriptional evidence of ISR activation was found in adipose tissue of people with *MFN2^R707W^*-related lipodystrophy^8^. We thus screened multiple tissues from the KI mice for ISR activation. mRNA levels of *Atf4*, *Atf5*, and *Chop,* all sentinel markers of the ISR, were increased in BAT and WAT of KI mice (**Figure 4A-C**) but were unchanged in liver, heart, and skeletal muscle (except for a modest rise in *Atf5*: WT 0.60 vs. KI 0.86 relative expression, p.adj = .03). Phosphorylation of eIF2α and protein expression of Mthfd2, an Atf4-upregulated enzyme playing a rate-limiting role in mitochondrial one carbon metabolism^47–49^, were also both strongly increased in BAT and in WAT in KI mice (**Figure 4D-F & Figure 4-figure supplement 1E-G**), whereas they were unchanged in the liver, skeletal muscle and heart. mRNA expression of two important secreted mediators of the organismal metabolic response to mitochondrial dysfunction, Gdf15 and Fgf21, trended towards being increased in adipose tissue, but serum concentrations were unchanged in KI mice (**Figure 4-figure supplement 1A-D**). This differs from observations in patients with MFN2^R707W^-related lipodystrophy^49–51^, and may relate to the fact that such patients manifest non-alcoholic fatty liver disease, which was not seen in the KI animals.

**Figure 4.**
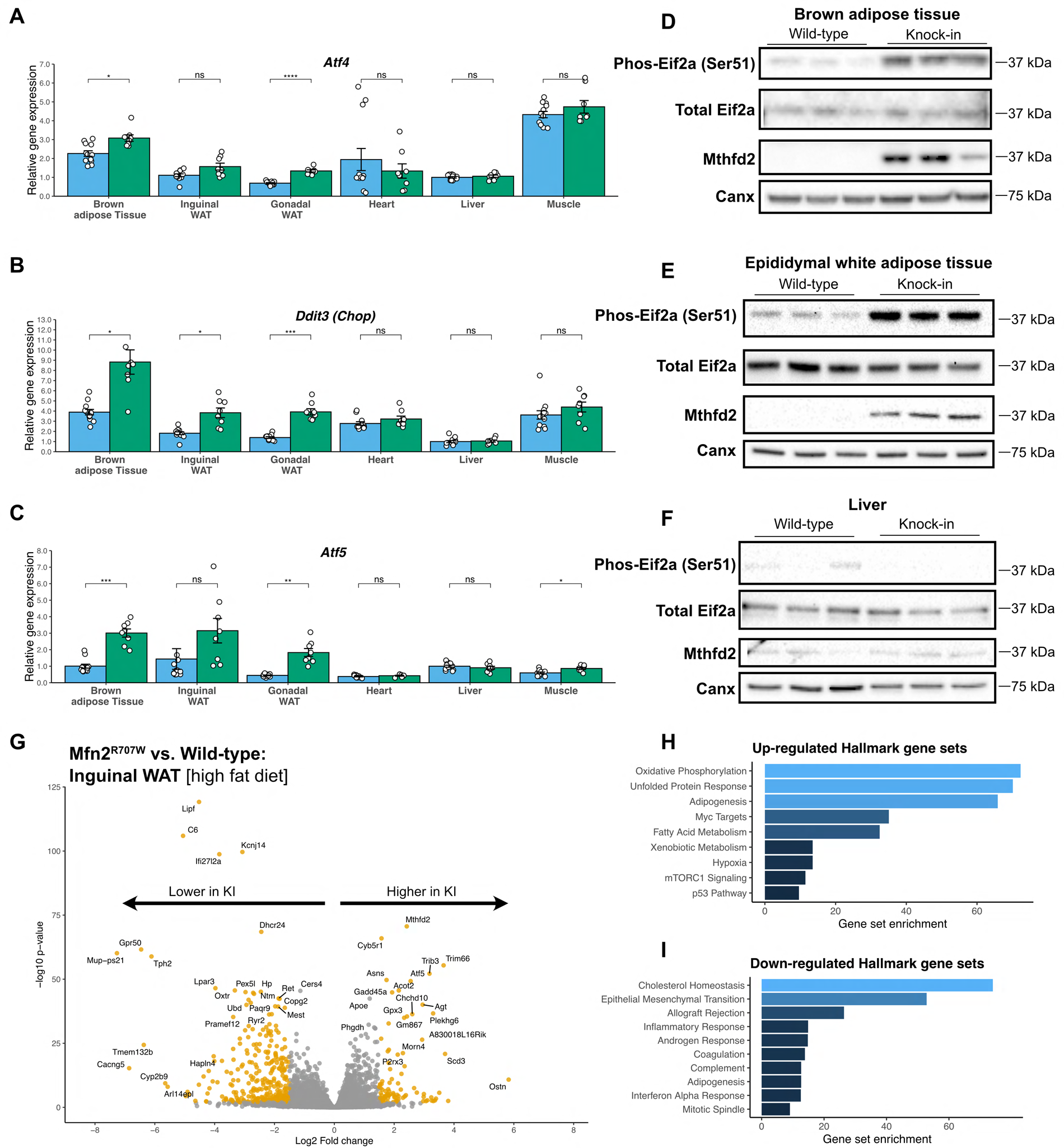
Mfn2^R707W^ causes an adipose tissue-specific induction of the integrated stress response. qPCR of genes involved in the integrated stress response (ISR) for six tissues from animals fed chow for 6 months: *Atf4* (A), *Ddit3* (Chop, B), and *Atf5* (C). Each data point represents one animal. Target gene CT values were normalised to three housekeeping genes (*36b4, B2m,* and *Hprt*) and expressed relative to WT liver for each gene. WT in blue, homozygous Mfn2^R707W^ in green. p-values are FDR-adjusted for multiple tests. Western blots from brown adipose tissue (BAT, D), epididymal WAT (E), and liver (F) illustrating Ser51-phosphorylation of eIF2a and expression of Mthfd2 with calnexin (*Canx*) as loading control. Western blots are representative of at least three biological and technical replicates. (G) Volcano plot from bulk RNA sequencing (n=8 per genotype) of inguinal WAT from mice on HFD. Significantly differentially expressed genes (Log2 fold change >1.5 and p-FDR <.001) are highlighted in orange. Pathway analysis using significantly differentially expressed genes for upregulated (H) and downregulated (I) Hallmark gene sets. All illustrated gene sets are enriched with p-FDR <.05. * p-FDR <.05, ** p-FDR <.01, *** p-FDR <.001. **Figure 4-source data.** Raw and annotated Western blots for Phospho-/Total-Eif2a, Mthfd2, and loading calnexin from 4D, 4E, and 4F.

To obtain an unbiased view of the transcriptional consequences of the lipodystrophy-associated Mfn2 mutation, we next applied bulk RNA sequencing (RNAseq) to BAT and inguinal WAT from high fat diet fed mice. Inguinal, rather than epididymal, WAT was selected as subcutaneous adipose tissue is predominantly affected in human MFN2-related and other partial lipodystrophies^52^. Induction of the ISR was confirmed in both BAT and WAT (**Figure 4G-H & Figure 4-figure supplement 2**), with the “unfolded protein response” gene set, a surrogate for the ISR, the top upregulated gene set in BAT and 6th in inguinal WAT. *Atf5* and *Mthfd2* were confirmed among the most highly upregulated mRNAs in both tissues, and a range of other well established ISR genes also showed increased expression. These included *Ddit3* (Chop), *Trib3*, an Atf4-driven ISR component that exerts negative feedback on the ISR, and *Gadd45a*, involved in ISR-induced G2/M checkpoint arrest^53^.

The top upregulated gene set in inguinal WAT was “oxidative phosphorylation”, driven solely by increased expression of nuclear-encoded mitochondrial genes (**Figure 4-figure supplement 1H**). In contrast, mitochondria-encoded genes were nearly universally downregulated (**Figure 4-figure supplement 1I**), recapitulating the pattern seen in affected human WAT^8^. In BAT, a similar but weaker pattern was seen on inspection of heatmaps, but this was not sufficient to reach statistical significance.

Another finding common to mouse and human was the transcriptional evidence of mTORC1 activation. The “mTorc signalling” gene set was the second most upregulated group in BAT and third most upregulated in WAT (**Figure 4-figure supplement 2B&E**). This activation is consistent with the proposed role for mTORC1 in mediating the proximal ISR^45^, and is of interest given accumulating evidence that the mTORC1 inhibitor sirolimus may exert beneficial effects in various mitochondrial diseases^54^.

Although no increase in adipose tissue mass was seen in *Mfn2^R707w^*, the “adipogenesis” gene set was upregulated in inguinal WAT under both diet conditions (**Figure 4H & Figure 4-figure supplement 2B**). However, closer inspection revealed a mixed profile of individual gene expression. The most consistent finding was the downregulation of the adipokine-encoding mature adipocyte genes *Adipoq* and *Lep* in *Mfn2^R707w^* (discussed below). *Adipoq* and *Lep* were also modestly downregulated in BAT (*Adipoq*: -0.65 log_2_ fold change (log_2_FC), p.adj = 6.3×10^-4^; *Lep*: -1.7 log_2_FC, p.adj = .03), but the adipogenesis gene set was not enriched overall (**Figure 4-figure supplement 2E**).

To assess for other potential drivers of adipose hyperplasia, we also examined downregulated gene sets. The signature of epithelial-mesenchymal transition (EMT) was the most strongly down-regulated set in mouse BAT and WAT (**Figure 4I & Figure 4-figure supplement 2C&F**), and was also previously found to be downregulated in overgrown human WAT in MFN2-associated MSL^8^. In bulk transcriptomic data it is not possible to discern the cell type(s) responsible for this consistent signature. However TGFβ family ligands are important mediators of EMT, some family members inhibit adipogenesis^55^, and they also play important roles in regulating mitochondrial function and in responding to mitochondrial dysfunction^56^.

We next sought to assess whether the increased demand for adipose expansion imposed by HFD feeding exacerbates the transcriptional consequences of Mfn2 R707W homozygosity. RNAseq was thus undertaken of inguinal WAT from mice maintained on HFD for 6 months. Comparison to WT animals revealed strikingly concordant findings to those seen in chow-fed mice (**Figure 4-figure supplement 2G**). No general differences were seen in the magnitude of transcriptional changes induced by Mfn2 R707W homozygosity between conditions. Oxidative phosphorylation, unfolded protein response and adipogenesis genes and gene sets were strongly upregulated, whereas genes in the EMT set were downregulated in KI mice on both diets. An exception was the group of mRNAs related to cholesterol homeostasis, for which diet strikingly modified the effect of genotype. They were the top downregulated gene set in inguinal WAT in HFD-fed animals but were not significantly altered on chow in the same depot and were upregulated in BAT (**Figure 4I & Figure 4-figure supplement 2E**). Lower expression of key enzymes in cholesterol metabolism (e.g. Hydroxymethylglutaryl-CoA synthase (*Hmgcs1*), mevalonate kinase (*Mvk*), and squalene monooxygenase (*Sqle*)) in WAT on HFD is consistent with the response to inhibition of the mitochondrial respiratory chain in primary human fibroblasts^57^.

### Lower circulating leptin and adiponectin in mice homozygous for Mfn2^R707W^

One of the most striking aspects of the Mfn2^R707W^-associated lipodystrophy phenotype is the low or undetectable serum leptin concentration despite abundant whole body adiposity, accounted for mostly by excess upper body adipose tissue of relatively normal histological appearance^8^. Serum adiponectin concentrations are also low, however this is in keeping with the “adiponectin paradox” widely seen in obesity with insulin resistance^58^. Mirroring these human observations, KI mice showed low serum leptin and adiponectin concentrations on both chow and HFD (**Figure 5A-C**), though unlike humans, the mice had normal fat mass and insulin sensitivity. In both WT and KI mice serum leptin concentrations correlated positively with whole body adiposity on chow and HFD, but a generalised linear model revealed marked attenuation of the relationship between serum leptin concentration and adipose mass in the KI mice (**Figure 5C & Figure 5 supplement 1A**).

**Figure 5.**
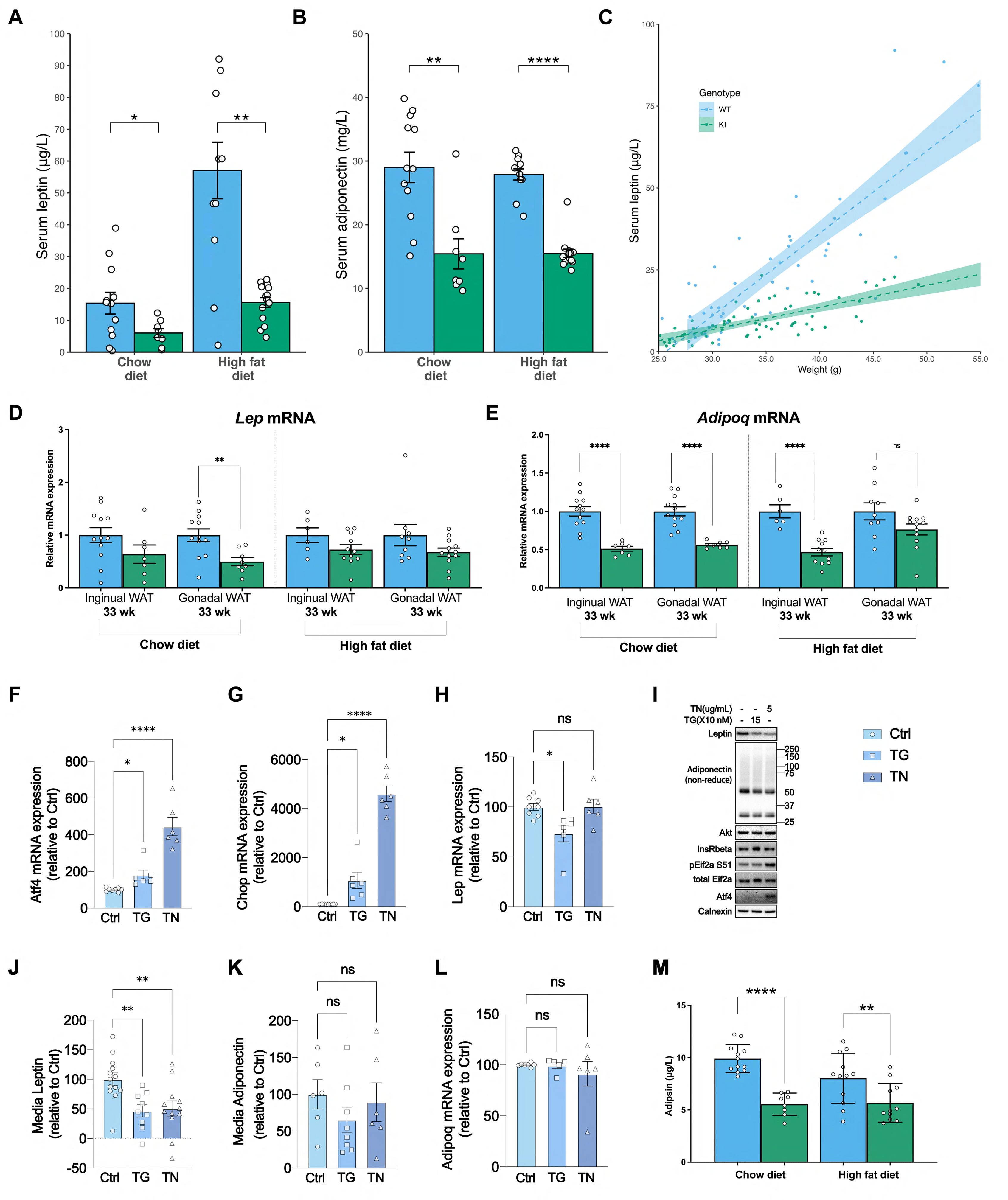
Mfn2^R707W^ decreases adipose secretion of leptin and adiponectin. Fasting serum leptin (A) and adiponectin (B) from mice after 6 months on chow or HFD. Asterisks indicate t-test comparing WT and KI with p-values adjusted for multiple testing. (C) Relationship between leptin and body weight for animals fed a HFD. Each data point represents one measurement of leptin, with multiple measurements per animal. Shaded area represents the 95% confidence interval. Data from n=13-14 animals. qPCR of leptin (D) and adiponectin (E) from white adipose tissue (WAT) depots. Each data point represents data from a separate animal. Target gene CT values were normalised against three housekeeping genes (*36b4, B2m,* and *Hprt*) and expressed relative to WT for each condition. (F-H) mRNA expression of *Atf4*, *Chop*, and *Lep* in primary adipocytes treated with either DMSO control or Thapsigargin (TG, 150 nM) or Tunicamycin (TN, 5 μg/mL) for 6 hr. Each data point represents an individual well from a separate biological experiment (n=5). Asterisks indicate significance on one-way ANOVA with correction for multiple comparisons. (I) Representative western blots of primary adipocytes treated with either DMSO control or Thapsigargin (TG, 150 nM) or Tunicamycin (TN, 5 μg/mL) for 6 hr. (J) Leptin and (K) Adiponectin secretion from primary adipocytes treated with either DMSO control or Thapsigargin (TG, 150 nM) or Tunicamycin (TN, 5 μg/mL) over 6 hr. Asterisks indicate significance on one-way ANOVA with correction for multiple comparisons. (L) mRNA expression of *Adipoq* in primary adipocytes treated with either DMSO control or Thapsigargin (TG, 150 nM) or Tunicamycin (TN, 5 μg/mL) for 6 hr. (M) Fasting serum adipsin (complement factor D) from mice after 6 months on chow or HFD. * p-FDR <.05, ** p-FDR <.01, *** p-FDR <.001, **** p-FDR <.0001. **Figure 5-source data.** Raw and annotated Western blots for all proteins from panel 5I.

RNAseq suggested that *Lep* mRNA in inguinal WAT was significantly lower in Mfn2^R707W^ compared to WT mice under both chow (-1.5 log_2_ fold change (log_2_FC), p.adj = 3.2×10^-4^) and HFD (-0.6 log_2_FC, p.adj = 5.3×10^-4^). *Adipoq* mRNA was also significantly lower in chow (-0.6 log_2_FC, p.adj = 2.7×10^-4^) and HF fed mice (-0.8 log_2_FC, p.adj = 3.3×10^-15^). RT-qPCR analysis confirmed lower *Lep* mRNA in epididymal WAT from chow fed Mfn2^R707W^ animals (**Figure 5D**). RT-qPCR analysis also found 50% lower *Adipoq* mRNA across both inguinal and epididymal WAT in chow fed Mfn2^R707W^ mice and in inguinal WAT from HFD fed Mfn2^R707W^ mice (**Figure 5E**).

To assess leptin secretion directly, we studied production of adipokines from adipose explants. Explants from KI mice fed on HFD for 4 weeks showed lower secretion of leptin and adiponectin per gram of tissue (**Figure 5-figure supplement 1A-B**). KI explants also exhibited minimal increase in leptin secretion after insulin and dexamethasone stimulation. *Adipoq* mRNA was lower in KI than WT explants at baseline whereas *Lep* mRNA was no different at baseline but failed to increase in KI explants following insulin and dexamethasone stimulation (**Figure 5-figure supplement 1C**).

To assess whether induction of the ISR in adipose tissue may be responsible for the relative leptin deficiency in both humans and mice homozygous for MFN2 R707W, we studied adipocytes freshly isolated from mouse gonadal fat in floating culture, adapting a recently described protocol^59^. We induced the ISR using each of two different well characterised activators, namely thapsigargin (TG), an inhibitor of the endoplasmic reticulum (ER) Ca^2+^ ATPase that depletes ER calcium^60, 61^, or tunicamycin (TN), which blocks protein glycosylation. ISR induction was confirmed by increased *Atf4* and *Ddit3* mRNA and/or protein expression, and by eIF2α phosphorylation (**Figure 5F-I**). *Lep* mRNA was modestly reduced by TG but not by TN (**Figure 5H**), whereas both agents reduced intracellular leptin protein expression (**Figure 5I**) and secretion (**Figure 5J**). Expression of adiponectin was also reduced without a change in mRNA level whereas expression of Akt and the insulin receptor beta subunit were not altered (**Figure 5I, K & L**). These data suggest that ISR activation may have a bigger impact on secreted proteins than on intracellular proteins, in keeping with the previous suggestion that the ISR tends to prevent a fall in intracellular amino acid concentrations^61^. In seeking to validate this notion, we proceeded to measure the serum concentration of adipsin, another ‘adipokine’ selectively secreted by adipocytes. Serum adipsin concentrations were significantly lower in the KI mice than in WT controls in both chow and HFD fed mice (**Figure 5M**), suggesting that low serum leptin and adiponectin may be part of a wider pattern of reduced adipocyte-derived secreted proteins in *MFN2^R707W^* KI mice.

## Discussion

The recent discovery that humans homozygous for the MFN2 R707W mutation manifest striking adipose redistribution associated with serious metabolic disease is probably the clearest example to date of a causal link in humans between a mitochondrial perturbation and adipose dysregulation. MFN2-related lipodystrophy has some remarkable and currently poorly understood features. These include: a) a marked and often dramatic increase in upper body adiposity, contrasting with loss of lower limb adipose tissue; b) a severe reduction in plasma leptin concentration despite abundant, histologically near-normal upper body fat. These problems have to date been associated only with the R707W mutation. To investigate the molecular pathogenesis of MFN2 R707W-related lipodystrophy, and the role of MFN2 in leptin synthesis and secretion, we generated and characterised homozygous *Mfn2^R707W/R707W^* mice.

*Mfn2* knock-out mice die in early embryogenesis^30^ while mice homozygous for either of two human neuropathy-associated, GTPase null missense mutations (H361Y or R94W) die on day 0-1^32^. Mfn2 is nearly ubiquitously co-expressed with its closely related paralogue Mfn1, and this demonstrates that it has essential, non-redundant functions. Homozygous Mfn2 R707W mice, in contrast, were viable and bred normally, showing that Mfn2 R707W retains significant function. Mfn2 also has key metabolic functions in mature adipocytes: mice lacking Mfn2 in all adipocytes^35^ or in brown adipocytes alone^36^ did not show reduced adipose tissue, but they did exhibit lower energy expenditure, reduced expression of multiple oxidative phosphorylation subunits, and impaired cold tolerance. Paradoxically, however, both lines were protected from systemic insulin resistance. Mice in which Mfn2 was deleted in all adipocytes in adulthood showed increased obesity and elevated blood glucose^37^. In contrast, homozygous Mfn2 R707W mice showed no overt change in adipose mass, metabolic function, or thermogenic capacity even though the genetic alteration was constitutional. This confirms some retained Mfn2 function also in adipose lineages.

Primary anatomical and/or functional defects in humans with MFN2 R707W homozygosity have been observed only in adipose tissue and peripheral nerves, with some but not all people reported to have sensorimotor neuropathy. Such neuropathy is commonly observed in people with heterozygous loss of MFN2 function^14, 62^. Although homozygous Mfn2 R707W KI mice had no overt anatomical adipose abnormality, and failed to show obvious neurological phenotypes, ultrastructural studies did reveal mitochondrial network disruption in adipose tissues, but not liver, skeletal muscle, or heart. The structural changes in mouse adipose mitochondria resembled those in adipose tissue from patients homozygous for the MFN2 R707W mutation^8^, and in both species these were associated with robust activation of the integrated stress response (ISR), which was also seen previously in tissue-specific *Mfn2* knock-out mice^63^. The ISR was not activated in liver, muscle, and heart, strengthening evidence that Mfn2 R707W has deleterious effects selectively in adipose tissue. We cannot conclusively exclude the possibility that tissues other than brown and white adipose tissue are affected, as we have not studied every tissue in the mice or patients, but if present, it is not associated with overt phenotypes.

The reason for adipose-selectivity of abnormalities in Mfn2 R707W homozygous mice is not established, but disruption of the function of Mfn2 in establishment or maintenance of mitochondrial-lipid droplet contact sites, perhaps through interaction with an adipose-specific protein, is plausible. We did observe reduced mitochondrial-lipid droplet contacts in brown adipose tissue from Mfn2 R707W KI mice, as reported in adipocyte *Mfn2* knock-out animals, but we were unable to replicate the direct mitofusin 2-perilipin 1 interaction previously reported using co-immunoprecipitation^35^. This requires further characterisation in the context of _Mfn2R707W._

Whether mitochondrial structural and functional perturbation mediates the overgrowth of some adipose depots and loss of others in humans with MFN2 R707W homozygosity remains to be proven. If it does, the mechanisms transducing dysfunction of a key organelle into cellular hyperplasia in some adipose depots but loss of adipose tissue in others are also unexplained. KI mice exhibit neither adipose loss nor hyperplasia, even when challenged by a HFD. This failure to model the gross anatomical adipose abnormalities of humans, despite evidence of mitochondrial dysfunction and attendant ISR establishes that the cellular abnormalities we describe are not sufficient to perturb adipose growth, but they may still be necessary. Whether a permissive genetic background, or an undefined additional stressor, are required as cofactors, remains to be determined.

Some of the transcriptomic changes observed that are common to mouse and human adipose tissue do suggest both potential opportunities to intervene pharmacologically, and mechanistic hypotheses relating to adipose hyperplasia that warrant further investigation. For example, transcriptional evidence of strong mTORC1 activation, likely part of the proximal ISR triggered by mitochondrial dysfunction, suggests that mTOR inhibitors such as sirolimus are worthy of testing. It is possible that they may restrain the ISR, thereby restraining compensatory adipose hyperplasia or even inducing synthetic lethality in cases of MFN2 R707W homozygosity. Several previous studies have suggested that mTOR inhibition may have beneficial effects in other primary mitochondrial disorders^54, 64^. Downregulation of TGFβ also merits testing as one candidate mechanism linking Mfn2 R707W homozygosity to adipose hyperplasia. This is based on strong transcriptional evidence of downregulated EMT in both mice and humans, on the important roles of TGFβ in EMT and adipogenesis^65, 66^, and on the inter-relationship of TGFβ signalling with mitochondrial dysfunction^67–70^.

A further notable difference between the mice and humans with the homozygous MFN2 R707W mutation is that serum concentrations of GDF15 and FGF21 were increased in people but not mice. This likely reflects the fact that affected humans also have fatty liver disease and diabetes, both strongly associated with elevated stress hormone levels^71, 72^. In keeping with this, we have shown in mice that the liver is the predominant source of circulating FGF21 and GDF15, with little or no contribution from adipose tissue^73^.

Although abnormal adipose growth and metabolic disease were not seen in KI mice, the low plasma leptin concentration seen in human adipose overgrowth associated with MFN2 mutations was replicated. Leptin concentrations were not reported in previously described adipocyte *Mfn2* knock-out mice^35, 37^, and although a different model of adipose-specific mitochondrial dysfunction (*Tfam* knock-out) did show reduced serum leptin, fat mass was also reduced compared to WT^74^. Lower adipose leptin secretion caused by Mfn2 R707W does not appear to be predominantly transcriptionally mediated in mice, as in some analyses, for example of adipose explants under basal conditions, leptin secretion was reduced (**Figure 5-figure supplement 1A**) without alteration of leptin mRNA (**Figure 5-figure supplement 1C**). Our findings suggest instead that the lower leptin secretion is a consequence of ISR activation. Synthesis of adipokines is an amino acid-intensive process, and activation of the UPR typically results in conservation of amino acids, in part through reduction of protein secretion^61, 75–77^. Stressing primary adipocytes with tunicamycin or thapsigargin reduced leptin secretion without any effect on expression of non-secreted proteins such as the insulin receptor. We also observed upregulation of pathways related to amino acid metabolism (particularly in BAT) (**Figure 4 – Figure Supplement 2**), which would be consistent with the known transcriptional effects of Atf4^78^. This suggests that the low leptin may not be due to a mitofusin-specific mechanism, rather secondary to activation of the ISR in adipose tissue as part of amino acid conservation. KI mouse data suggest that other adipokines, including adiponectin and adipsin, are similarly affected.

This study has limitations. We characterised male homozygous KI mice only in detail, so cannot extrapolate our results to females with confidence, though case series do not suggest significant sexual dimorphism in the human disorder^8–10^. We also did not study heterozygous animals, but as human MFN2 R707W-associated lipodystrophy shows recessive inheritance, and as even homozygous mice do not exhibit lipodystrophy, a phenotype in heterozygous animals seems unlikely. It is unclear to what extent the phenotype observed in BAT is due to reduction in expression of both the mitofusins. Given the concomitant reduction in expression of Oxphos components in BAT, the lower mitofusin expression may be more reflective of general mitochondrial perturbation. Lastly, this study has not directly assessed the ability of Mfn2^R707W^ mutants to mediate mitochondrial fusion. However given the normal mitochondrial network morphology in non-adipose tissues in homozygous KI mice and in dermal fibroblasts from humans homozygous for MFN2 R707W^8^, any defect is likely mild and context dependent.

## Conclusion

Mfn2^R707W^ KI mice show adipose-selective alteration of mitochondrial morphology and robust activation of the integrated stress response, but no abnormal adipose growth or systemic metabolic derangement. The KI mice do show suppressed leptin expression and plasma leptin concentration, likely secondary to the adipose selective mitochondrial stress response. The unique association of human lipodystrophy with the MFN2 R707W allele remains unexplained, but transcriptomic analysis suggested that reduced TGFβ signalling warrants further explanation as a potential cause of adipose hyperplasia, while mTOR inhibitors are worthy of testing in models as a potential targeted therapy.

## Methods

### Generation of the *Mfn2^R707W/R707W^* knock-in mouse

*Mfn2^R707W^* mice were generated using CRISPR-Cas9 microinjection of fertilised oocytes at The Wellcome Trust Centre for Human Genetics (Oxford, UK). Two single guide RNA (sgRNA) sequences targetting exon 18 of the mouse *Mfn2* gene (ENSMUSE00000184630) were used, namely 5’-CACCgTTCCTGCTCCAGATTATCTC-3‘ and 5‘-AAACGAGATAATCTGGAGCAGGAAc-3‘. The single-strand donor oligonucleotide (ssODN) incorporated the desired R707W mutation by recoding codon 707 from CGA to TGG. (This involved two point mutations in order to avoid a stop codon.) A silent mutation was added upstream to generate an EcoRV restriction site (Figure 1).

Superovulated 3-week old C57BL/6J female mice were mated with C57BL/6J males. Embryos were extracted on day 0.5 of pregnancy and cultivated until two pronuclei were visible. One pronucleus was injected with purified sgRNA (20ng/μL), Cas9 protein (100 ng/μL), and the ssODN template (10ng/μL). Embryos were reimplanted into pseudopregnant CD1 foster mothers at day 0.5 post-coitum. Mfn2^R707W^ was confirmed by Sanger sequencing of F0 founder males (with one additional upstream silent mutation ACC>ACA). Following cryopreservation of embryos, the line was re-derived in a colony of C57BL/6J mice in Cambridge, UK. Genotyping utilised the upstream EcoRV restriction enzyme digestion site. For genotyping, ear biopsies were digested in Chelix and proteinase K (0.1mg/mL) for 45 min at 50°C to extract gDNA for use in PCR. gDNA was amplified using primers 5’-AGTCCCTTCCTTGTCACTTAGT-3’ and 5’-ATCTCACAAGAAAGCGAAATCC-3’ and GoTaq DNA Polymerase (Promega), then digested using EcoRV (New England Biosciences). Wild-type (WT) *Mfn2* generates a PCR product of 523bp, homozygous knock-ins (KI) have two bands at 327bp and 196bp, and heterozygotes have all three bands (Figure 1).

### Mouse husbandry and phenotyping

All experiments were performed under UK Home Office-approved Project License 70/8955 except for thermogenic capacity assessments which were conducted under P0101ED1D. Protocols were approved by the University of Cambridge Animal Welfare and Ethical Review Board. Animals were co-housed in groups of 2-5 littermates of mixed genotype, on 12 hr light/dark cycles. They had access to food and water *ad libitum* except when fasting prior to experimental procedures.

Male mice were used for all studies. WT and KI male mice aged 5 weeks were randomly allocated to HFD (45% kcal as fat, 4.7kcal/g, Research Diets D12451i) or chow (Safe Diets R105-25) for up to 6 months. Investigators were blinded to animal genotype at the point of data collection. Mice were weighed weekly. Tail blood samples were collected four weekly into heparinised capillary tubes (Hawksley) and spun at 13,000g for 4 min for plasma analysis of leptin and adiponectin. 6 hr fasted blood samples were obtained on weeks 16 and 24 of diet, and prior to sacrifice, for analysis of glucose, insulin, and lactate, whilst other blood samples were from fed animals. For lactate, tail vein blood was collected into fluoride oxalate tubes and centrifuged immediately before freezing of plasma at -80°C.

An intraperitoneal glucose tolerance test (IPGTT) was performed on week 31 of chow and HFD and an intraperitoneal insulin tolerance test (IPITT) was performed on week 32 of HFD (only) after a 6 hr fast. 1g/kg of glucose and 0.75 units/kg of insulin were administered for the IPGTT and IPITT, respectively. Blood glucose was measured at 0, 10, 20, 30, 60, 90, and 120 min after the injections using a glucometer (Abbot Laboratories) and glucose test strips (Abbot Laboratories). Insulin was measured at 0 min at the start of IPGTT from a tail vein blood sample.

Body composition (lean and fat mass) was assessed prior to sacrifice by Time-Domain Nuclear Magnetic Resonance (TD-NMR) using a Minispec Live Mouse Analyzer (Bruker). Mice were sacrificed at 4- or 33- weeks on diet after a 6 hr fast. Tissues were weighed, sections were removed for histological or electron microscopic analysis, and remaining tissue was snap frozen in liquid nitrogen.

To estimate the number of mice required for experimental groups, We used data from the adiponectin-Cre *Mfn2* knock-out mouse^35^, aiming to determine a difference between fat mass in Mfn2^R707W^ and wild-type. Mean fat mass in adipose-specific *Mfn2* knock-out = 3.8±0.43g. Mean fat mass in wild-type = 2.9±0.14g. For 80% power at 0.05 significance, to detect 0.9g difference, sample size: 8 animals per group.

### Calorimetry studies

Eight-week-old chow-fed male mice were housed in either cold (10°C) or thermoneutrality (30°C) for 4 weeks. Animals were anaesthetised using 90mg/kg of pentobarbital by intraperitoneal injection and were placed in 2.7 l calorimetry chambers (Oxymax, Columbus instruments, Ohio) attached to a Promethion calorimetry system (Sable Systems, Las Vegas, NV, USA) pre-warmed to 30°C for 20 min for measurement of basal energy expenditure. They were then given a subcutaneous injection of 2 µl/g of 0.5 mg/mL noradrenaline bitartrate (NA) plus 1.66 µl/g of 18 µg/µl pentobarbital and NA-stimulated energy expenditure was measured for 25 min. Animals were then sacrificed, and tissues snap frozen as described above. Basal energy expenditure was calculated from the three readings prior to NA-injection. Peak energy expenditure was calculated from the three greatest readings 5-25 min after NA-injection. NA-induced energy expenditure was calculated as the difference between peak and basal energy expenditure.

### Transmission Electron Microscopy (TEM)

Chow-fed mice aged 8 weeks were sacrificed and white adipose tissue (inguinal and epididymal), brown adipose tissue, skeletal muscle (quadriceps), heart, and liver were removed, cut into <1 mm^3^ cubes and fixed (2% glutaraldehyde/2% formaldehyde in 0.05 M sodium cacodylate buffer pH 7.4 containing 2 mM calcium chloride) on a rocker at 4℃ overnight. Samples were then washed five times with 0.05 M sodium cacodylate buffer pH 7.4 and osmicated (1% osmium tetroxide, 1.5 % potassium ferricyanide, 0.05 M sodium cacodylate buffer pH 7.4) for 3 days at 4°C.

Following initial osmication, samples were washed five times in DIW (deionised water) then treated with 0.1 % (w/v) thiocarbohydrazide/DIW for 20 min at room temperature in the dark. After washing five times in DIW, samples were osmicated a second time for 1 hr at room temperature (2% osmium tetroxide/DIW). After washing five times in DIW, samples were block stained with uranyl acetate (2% uranyl acetate in 0.05 M maleate buffer pH 5.5) for 3 days at 4°C. Samples were washed five times in DIW and then dehydrated in a graded series of ethanol (50%/70%/95%/100%/100% dry) 100% dry acetone and 100% dry acetonitrile, three times in each for at least 5min. Samples were infiltrated with a 50/50 mixture of 100% dry acetonitrile/Quetol resin (without benzyldimethylamine (BDMA)) overnight, followed by 3 days in 100% Quetol (without BDMA). Then, samples were infiltrated for 5 days in 100% Quetol resin with BDMA, exchanging the resin each day. The Quetol resin mixture is: 12 g Quetol 651, 15.7 g NSA, 5.7 g MNA and 0.5 g BDMA (all from TAAB). Samples were placed in embedding moulds and cured at 60°C for 3 days.

Thin sections were cut using an ultramicrotome (Leica Ultracut E) and placed on bare 300 mesh copper TEM grids. Samples were imaged in a Tecnai G2 TEM (FEI/Thermo Fisher Scientific) run at 200 keV using a 20 µm objective aperture to improve contrast. Images were acquired using an ORCA HR high resolution CCD camera (Advanced Microscopy Techniques Corp, Danvers USA).

Analysis was performed by manual measurement of individual mitochondria from all obtained images using Fiji^79^/ImageJ measurement tools by an investigator who was blinded to the genotype of tissues/cells. In all tissues, mitochondrial perimeter and aspect ratio (length/width) were determined. In addition, in brown adipose tissue, the number of cristae per mitochondrion and mitochondrial-lipid droplet contacts were quantified. There were insufficient mitochondrial-lipid droplet contact sites with high quality preservation to permit this in white adipose tissue. In liver, mitochondrial-endoplasmic reticulum contact sites were quantified. It was not possible to assess for mitochondria-ER contacts in other tissues due to quality of preservation.

### Isolation of mitochondria for Oroboros analysis

For *ex vivo* measurement of mitochondrial respiratory capacity, mitochondria were isolated from BAT and liver of 12 week old chow-fed male mice using the protocols from McLaughlin *et al*.^80^ for BAT and Fernández-Vizarra *et al.*^81^ for liver. In brief, tissues were isolated and washed in buffer ‘B’ then homogenised using a drill-driven Teflon pestle and borosilicate glass vessel. Homogenates were centrifuged at 800 x g for 10 min at 4°C to remove cellular debris. Supernatant was removed, and then re-centrifuged at 10,000 x g for 10 min at 4°C to enrich mitochondria. Supernatant was discarded and mitochondria were resuspended in buffer ‘A’. Mitochondria were quantified using a BioRad Protein Assay.

Fifty µg of protein were used per chamber for high-resolution respirometry, using Oroboros (Innsbruck, Austria). There was sequential injection of: 20 µl of 1 M glutamate and 10 µl of 1 M malate (liver only), 20 µl of 0.5 M ADP, 20 µl of 1 M pyruvate, 20 µl of 2 M succinate, 1 µl of 1 mM CCCP (liver only), 2 µl of 2 mM rotenone, and 2 µl of 2 mM antimycin A.

### mtDNA content assay

Relative mtDNA content was assayed using real-time quantitative polymerase chain reaction (RT-qPCR) quantification of mitochondrial *Rnr2* and nuclear *Hk2* DNA. DNA was extracted from snap frozen murine tissue, using a DNeasy kit (Qiagen) as per the manufacturer’s instructions. DNA was quantified on Nanodrop and diluted to 4 ng/ul. RT-qPCR was performed in triplicate for each sample using 8 ng DNA with primers for *Hk2* and *Rnr2*. mt*Rnr2*/n*Hk2* was calculated using the standard curve method and expressed relative to WT.

### Protein extraction for WB

Fifty mg of frozen tissue was crushed using a pestle and mortar in liquid nitrogen. Powdered tissue was dissolved in 800 µl of RIPA buffer (Sigma, R0278) containing protease (Sigma, 11836170001) and phosphatase inhibitors (Roche, 04906837001). Samples were sonicated twice for 5 s at 30 Hz before centrifugation at 10,000 x g for 5 min at 4°C. Supernatant was extracted and, for adipose samples, re-centrifuged to remove excess lipid.

Thirty-45 μg of protein lysates were mixed with NuPAGE 4x LDS buffer (ThermoFisher Scientific), containing 0.05% 2-mercaptoethanol, and denatured for 5 min at 95°C. Samples were run on 4-12% Bis-Tris gels (Invitrogen) and transferred onto a nitrocellulose membrane using iBlot-2 (ThermoFisher Scientific). Membranes were washed in Tris-buffered saline with 0.1% (vol/vol) Tween 20 (TBST, Sigma) before blocking in 5% (wt/vol) skimmed milk powder dissolved in TBST. Membranes were incubated with primary antibodies at 4°C for 16 hr, washed with TBST five times for 5 min, followed by incubation with horseradish peroxidase (HRP)-conjugated secondary antibodies for 1 hr at room temperature. Blots were developed using Immobilon Western Chemiluminescent HRP Substrate (Millipore) with images acquired on BioRad ChemiDoc Imaging system.

### RNA isolation and qPCR analysis

At the end of the study, tissues were harvested and immediately snap frozen in liquid nitrogen and stored at -80°C. For RNA isolation, 30-50 mg of tissue was placed in Lysing Matrix D tubes and homogenized in 800 µl TRI Reagent (T9424, Sigma) using the Fastprep-24 Homogenizer for 30 s at 4-6 m/s (MP Biomedical). Homogenate was transferred to an RNase free tube and 200 µl chloroform (Sigma) added. The samples were vortexed and centrifuged at 13,000 rpm for 15 min at 4°C. The upper phase was then transferred to an RNase free tube and mixed with an equal volume of 70% ethanol before loading onto RNA isolation spin columns. RNA was extracted using a RNeasy Mini Kit (74106, Qiagen) isolation kit following the manufacturer’s instructions.

Total RNA of 600 ng was quantified using Nanodrop and converted to cDNA using MMLV Reverse Transcriptase with random primers and RNase inhibitor (Promega). RT-qPCR was performed using SYBR Green or TaqMan Universal PCR MasterMixes (Applied Biosystems) on QuantStudio 7 Flex Real time PCR system (Applied Biosystems). Reactions were performed in triplicate and RNA expression was normalised to *36b4*, *Hprt*, and *B2m* expression using the standard curve method.

### Immunoassays

Mouse sera and plasma were analysed by the Cambridge Biochemical Assay Laboratory, University of Cambridge. Leptin was measured using a 2-plex Mouse Metabolic immunoassay kit from Meso Scale Discovery Kit (Rockville, MD, USA) according to the manufacturers’ instructions and with supplied calibrants. GDF15 was measured using a modified Mouse GDF15 DuoSet ELISA (R&D Systems) as an electrochemiluminescence assay on the Meso Scale Discovery platform. Adiponectin (K152BYC-2, MSD) was analysed individually using the Meso Scale Discovery Kit (Rockville, MD, USA). NEFA were analysed using the Free Fatty Acid Kit (half-micro test) (11383175001, Roche) and TG was measured using an enzymatic assay (DF69A, Siemens Healthcare). Alanine aminotransferase (ALT, product code DF143), aspartate aminotransferase (AST, product code DF41A), and total cholesterol (product code DF27) were measured using automated enzymatic assays on the Siemens Dimension EXL analyzer. Mouse Fgf21 was measured using an enzyme-linked immunosorbent assay kit (R&D/ Biochne, cat no. MF2100).

### Histological processing

Fresh tissue was fixed in 10% formalin for 24 hr at room temperature immediately following sacrifice. Tissue was then embedded in paraffin and 4 μm sections were cut then baked overnight at 50°C. For haematoxylin & eosin (H&E) staining, slides were dewaxed in xylene for 5 min twice, then dehydrated in 100% ethanol for 2 min twice. Following a 3 min water wash, slides were stained with filtered Mayer’s haematoxylin (Pioneer Research Chemicals) for 7 min and blued in water for 4 min. Slides were then stained with 1% aqueous eosin (Pioneer Research Chemicals) for 4 min and briefly washed in water before dehydrating in 100% ethanol (1 min, twice) and cleared in xylene (2 min, twice) and mounting with Pertex. Slides were imaged using a Axio Scan Z1 slide scanner (Zeiss). Lipid droplet area and hepatic steatosis was quantified automatically using Halo software (Indica Labs).

### Transcriptomic profiling in white and brown adipose tissue

RNA was isolated from three tissues/dietary conditions for RNA sequencing: (1) inguinal WAT from chow fed animals (n=7 WT, n=6 KI, single technical replicates); (2) inguinal WAT from HFD-fed animals (n=8 WT, n=8 KI, two technical replicates per animal); and (3) BAT from HFD-fed animals (n=6 WT, n=6 KI, single technical replicate). RNA was quantified using Agilent 2100 Bioanalyzer (Agilent Technologies Inc) and only samples with RNA Integrity Number ≥8 were used for library preparation. cDNA libraries were made using Illumina TruSeq RNA sample kits and sequencing was performed on Illumina NovaSeq 6000 with paired-end 150 bp reads (Novogene, Cambridge, UK). Raw reads all passed quality control for Q_score_, error rate distribution, and AT/GC distribution.

Adapter sequences were removed from raw FASTQ files using cutadapt^82^ and aligned to *Mus musculus* reference genome (GRCm38) using STAR^83^. Binary alignment/map (BAM) files were sorted using samtools^84^ and counts were performed using featureCounts^85^. Differential gene expression (DGE) between WT and KI was performed using DESeq2^86^, where significance was considered as a Benjamini-Hochberg false-discovery rate (FDR) corrected p-value <.01. Pathway analysis was performed with the EnrichR package for R^87–89^ using significantly differentially expressed genes to determine enriched Hallmark^90^ and Kyoto Encyclopaedia of Genes and Genomes (KEGG)^91^ gene sets. Gene sets with FDR-corrected p-value <.05 were considered enriched. Figures were generated in R 4.0.2^92^ using packages pheatmap, ggplot2, and dplyr.

### Adipose explant experiments

Inguinal (subcutaneous) and epididymal (visceral) adipose tissue was harvested from 12-week-old male C57BL/6J mice fed either chow or HFD for 4 weeks (i.e. 8-12 weeks of age). Tissue was placed in Hanks’ Balanced Salt Solution (HBSS, H9269, Sigma) and kept on ice before cutting into 1-2 mm fragments. Approximately 100 mg fragments were incubated in a 12-well plate with M199 media ± 7 nM insulin (Actrapid, Novo Nordisk) and 25 nM dexamethasone (D4902, Sigma). After 24 hr incubation, media were collected, spun down at 5,000 x g and stored at -80°C until leptin and adiponectin assay as above. Explant tissues were weighed and snap frozen for RNA analysis.

### Primary adipocyte experiments

Mature white adipocytes were isolated from 12-20-week-old chow-fed male or female C57BL/6J or C57BL/6N mice as previously described^59, 93^ with modifications. Briefly, after dissection, gonadal adipose tissue was washed, minced finely, and dissociated to a single-cell suspension in Hanks’ Balanced Salt Solution containing 2.25% (w/v) BSA (Sigma, Cat. H9269) and 1 mg/mL collagenase (Sigma, C6885-1G) for approximately 15 min at 37°C in a shaking incubator. Digested material was diluted with 6x volume of high glucose DMEM media (Sigma, Cat. D6546) and connective tissue or undigested pieces were removed by passing through a 100 μm nylon mesh (Fisherbrand, 11517532). Floating adipocytes were washed twice with 6x volume of high glucose DMEM media before being used for downstream experiments.

Mature adipocytes with a packed volume of 60 μL were then cultured in 500 μL of high glucose DMEM media (supplemented with 10% FBS (Gibco Cat. 10270-106), 2 mM L-glutamine (Sigma Cat. G7513) and 1 x Penn/Strep (Sigma Cat. P0781)) per well in the presence of 100 nM insulin in a 24-well plate. On the third day of in vitro culture, 100 μL of media was sampled followed by media replenishment. Cells were treated with indicated concentration of Thapsigargin or Tunicamycin for 6 hr. Media were sampled at the end of treatment and cells collected for subsequent RT-qPCR or western blotting. Adipokine secretion from adipocytes during the 6 hr window was calculated as X=B-A*400/500 (A/B = medium adipokine concentration before/after treatment).

### Statistical analysis

Continuous data were expressed as mean ± standard error (SE). Normally distributed data were analysed by t-test (for two group pairwise comparison) and one-way ANOVA (for three or more groups) with post-hoc Bonferroni multiple comparisons test. FDR-corrected p-value <.05 was considered significant.

Statistical tests of TEM data (pairwise comparisons of mutant to WT) were based on the mean of independent biological replicates (i.e. the number of different samples, not the number of mitochondria) and FDR adjustment for the number of tests^94^.

All experiments were conducted at least three times using, where possible, randomisation of sample order and blinding of experimenters handling samples. No data were excluded from analysis. Data were analysed using R 4.0.2^92^ and GraphPad Prism version 9 (GraphPad, San Diego). Figures were made using BioRender.

### Materials availability

All reagents used are publicly available. Primer sequences and antibodies are detailed in Supplementary Tables 1 and 2. Code used in analysis is available from: https://doi.org/10.5281/zenodo.5770057. Raw counts from transcriptomic analysis are available from GEO with accession number GSE210771.

**Table 1:**
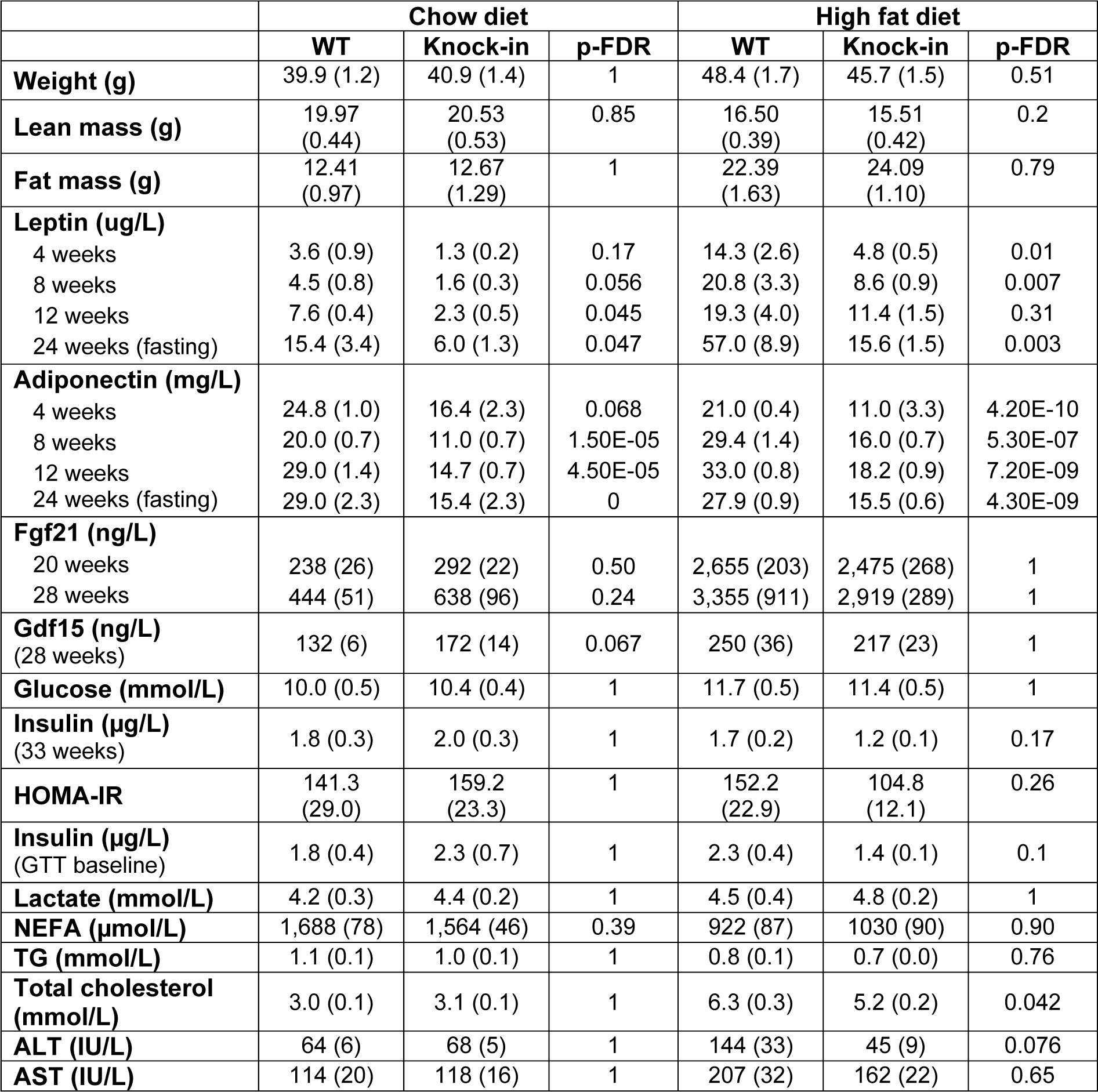
Weights and serum biochemistry for mice on chow or high fat diet for 6 months. Seven-week-old mice were fed chow (n=8-12) or 45% kcal HFD (n=13-14) for 6 months. Blood was taken 4 weekly with 6 hr fasting blood taken on week 28. p-FDR are false-discovery rate adjusted p-values derived from unpaired T-tests. Values in brackets referred to standard error of the mean. ALT, alanine aminotransferase; AST, aspartate aminotransferase; FGF21, fibroblast growth factor 21; Gdf15, Growth and differentiation factor 15; HOMA-IR, Homeostatic Model Assessment for Insulin Resistance; NEFA, non-esterified fatty acids.

**Table 2:**
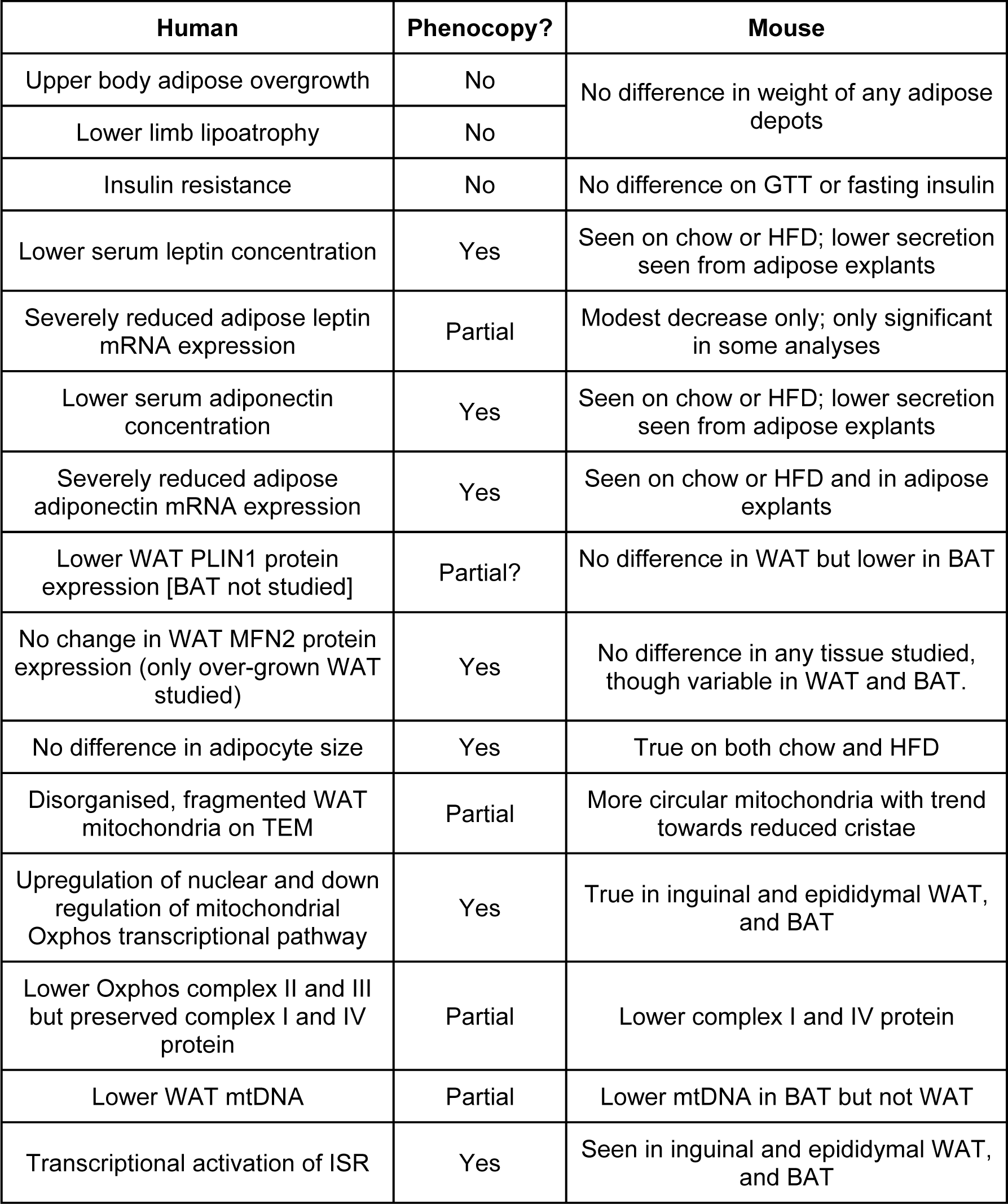
Comparison of human *MFN2^R707W^*-associated lipodystrophy with phenotype of *Mfn2^R707W/R7007W^* mice. BAT, brown adipose tissue; GTT, glucose tolerance test; HFD, high fat diet; ISR, integrated stress response; MFN2, mitofusin 2; mRNA, messenger ribose nucleic acid; mtDNA, mitochondrial DNA; PLIN1, perilipin 1; TEM, transmission electron microscope; WAT, white adipose tissue.

## Acknowledgements

We are grateful to the team at the University of Cambridge Advanced Imaging Centre for access to their facilities and for their expertise. We thank the Disease Model Core from the Wellcome-MRC Institute of Metabolic Science for their technical assistance in animal work. We also thank the Genomics and Transcriptomics core, the Histology core and G. Strachan from the Imaging core for technical assistance. Finally we thank M. Mimmack for his assistance with *in vitro* experiments.

## Figure Legends

**Figure 1-figure supplement 1.**
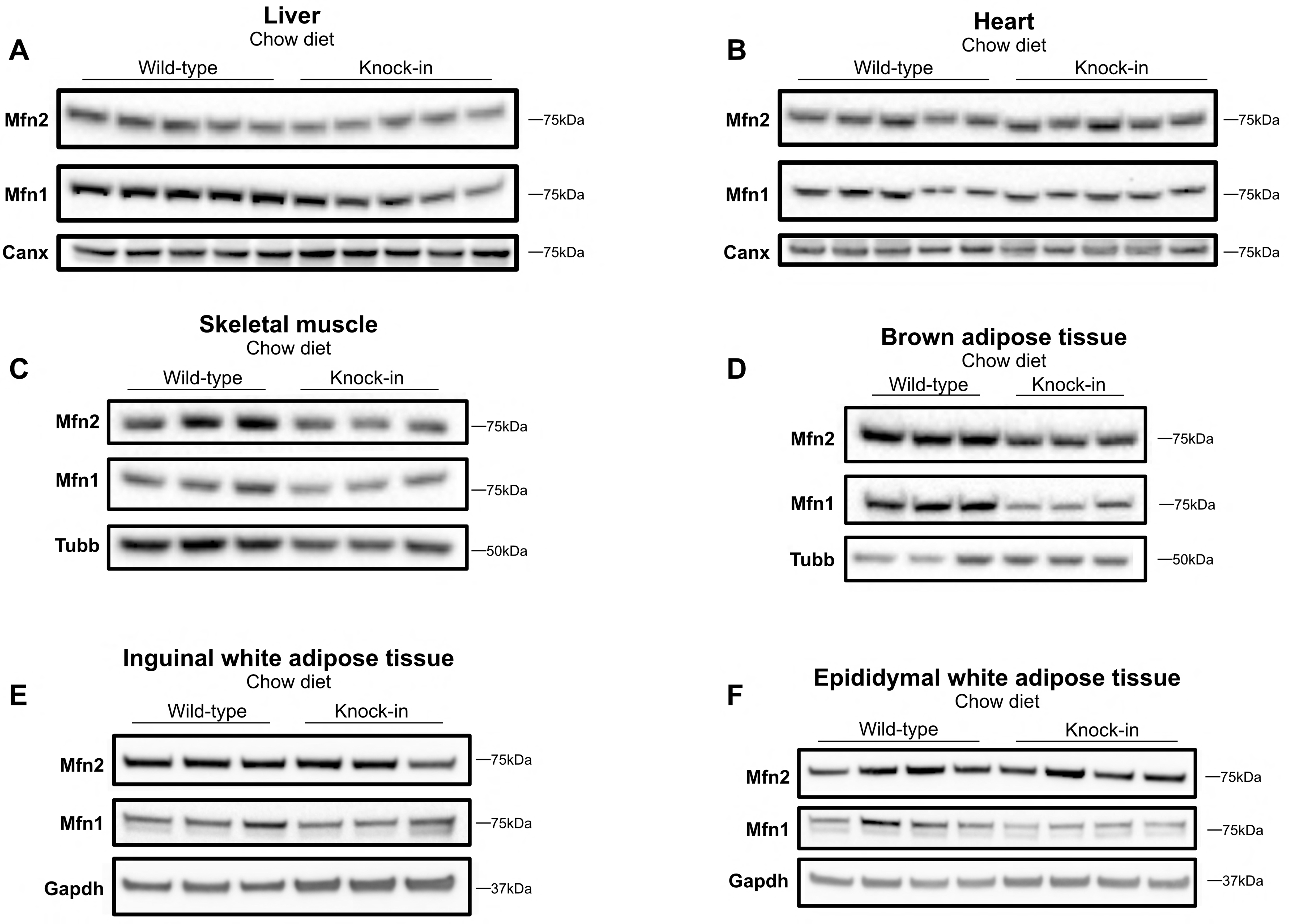
Expression of mitofusins on chow. Western blots showing expression of Mfn2 and Mfn1 in liver (A), heart (B), skeletal muscle (C), brown adipose tissue (D), inguinal white adipose tissue (E), and epididymal white adipose tissue (F) in mice fed chow for 6 months. Each lane contains samples from a separate animal and blots are representative of at least three replicates. *Canx* (calnexin), *Gapdh*, and *Tubb* (β-tubulin) are loading controls. **Figure 1-figure supplement 1-source data.** Raw and annotated Western blots for Mfn1, Mfn2, and loading control for all tissues.

**Figure 1-figure supplement 2.**
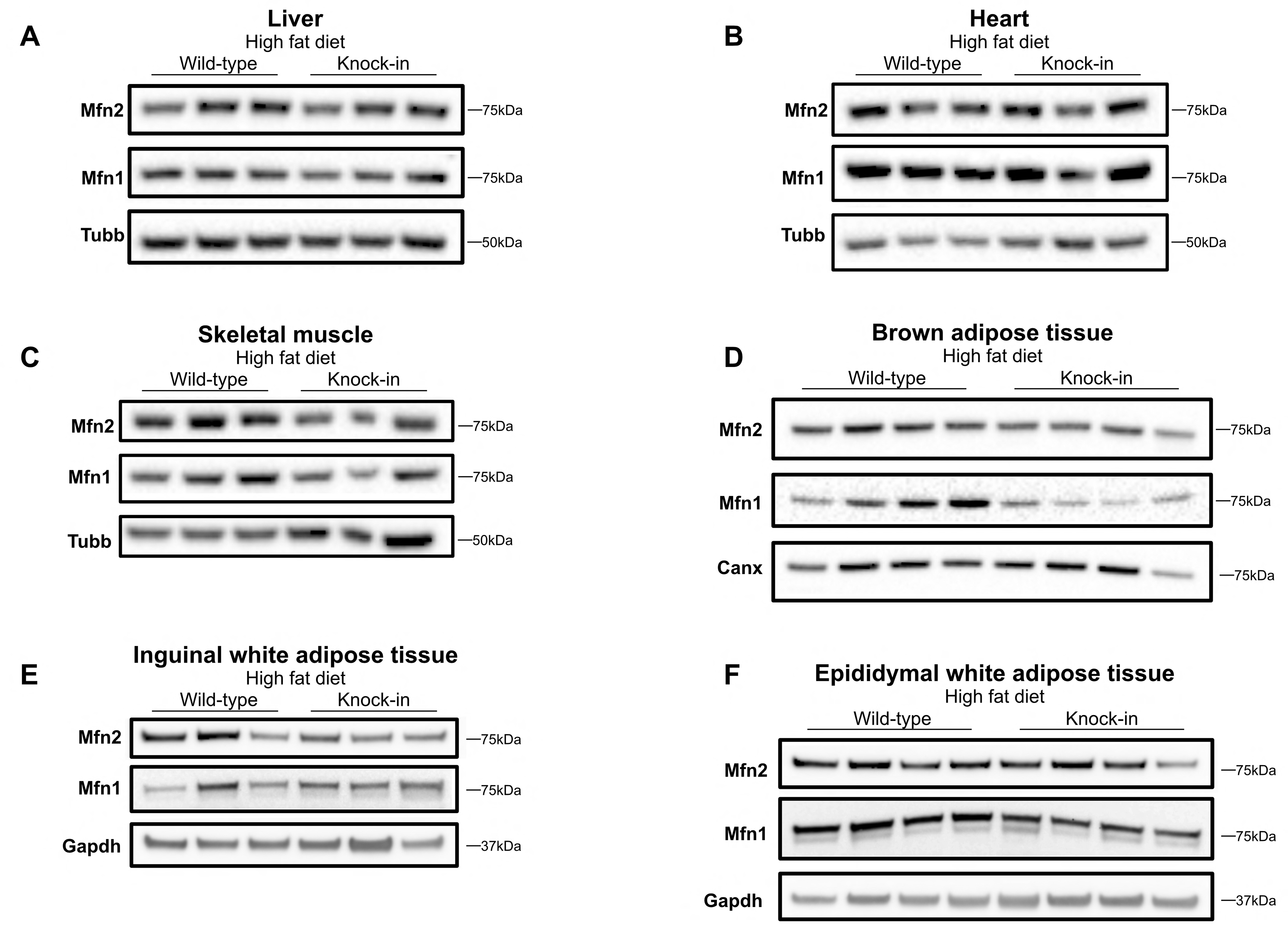
Expression of mitofusins on high fat diet. Western blots showing expression of Mfn2 and Mfn1 in liver (A), heart (B), skeletal muscle (C), brown adipose tissue (D), inguinal white adipose tissue (E), and epididymal white adipose tissue (F) of mice fed a 45% kcal HFD for 6 months. Each lane contains samples from a separate animal and blots are representative of at least three replicates. *Canx* (calnexin), *Gapdh*, and *Tubb* (β-tubulin) are given as loading controls. **Figure 1-figure supplement 2-source data.** Raw and annotated Western blots for Mfn1, Mfn2, and loading control for all tissues.

**Figure 2-figure supplement 1.**
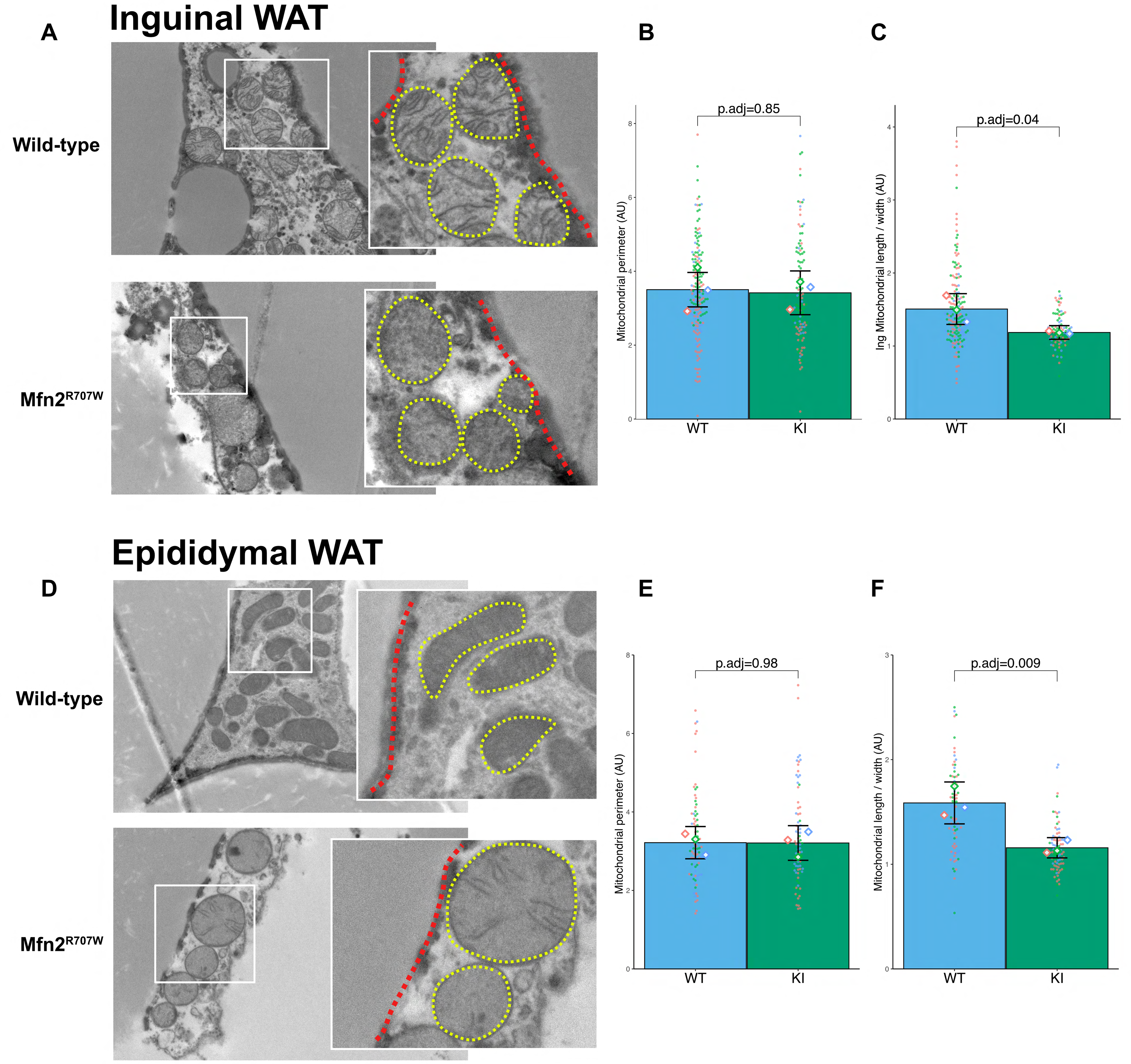
Effect of Mfn2^R707W^ on mitochondrial morphology in white adipose tissue. (A) Representative transmission electron microscopy (TEM) images from inguinal white adipose tissue (WAT) with zoomed-in images of mitochondria (highlighted in yellow) bordering lipid droplets (outlined in red). Quantification of mitochondrial perimeter (B) and aspect ratio (C) from TEM in inguinal WAT. Each dot represents an individual mitochondrion with diamonds showing biological replicates. p.adj gives the false-discovery rate (FDR) adjusted p-value from across all TEM analyses. (D) TEM images from epididymal WAT with zoomed-in images of mitochondria (highlighted in yellow) bordering lipid droplets (outlined in red). Quantification of mitochondrial perimeter (E) and elongation (F) from TEM on epididymal WAT. WT in blue, homozygous Mfn2^R707W^ in green.

**Figure 2-figure supplement 2.**
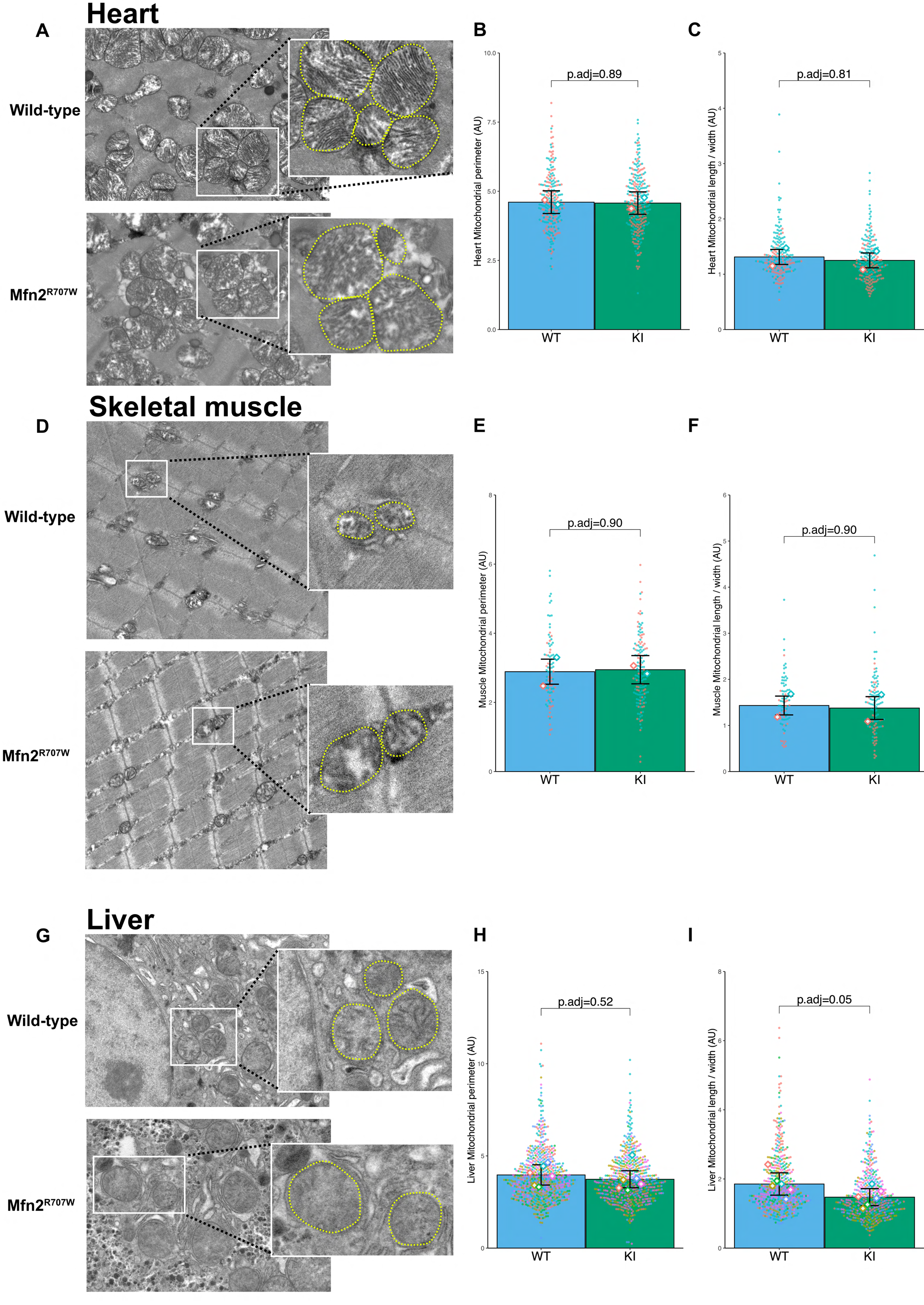
Effect of Mfn2^R707W^ on mitochondrial morphology in liver, heart, and muscle. (A) Representative transmisison electron microscopy (TEM) images from the heart with zoomed-in images of mitochondria (highlighted in yellow). (B) Quantification of mitochondrial perimeter from TEM on heart. Each dot represents an individual mitochondrion with each diamond showing biological replicates. p.adj gives the false-discovery rate (FDR) adjusted p-value from across all TEM analyses. (C) Quantification of mitochondrial elongation (length/width) from TEM on heart. (D) TEM images from skeletal muscle with zoomed-in images of mitochondria (highlighted in yellow). Quantification of mitochondrial perimeter (E) and elongation (F) from TEM on skeletal muscle. (G) TEM images from liver with zoomed-in images of mitochondria (highlighted in yellow). Quantification of mitochondrial perimeter (H) and elongation (I) from TEM on liver. WT in blue, homozygous Mfn2^R707W^ in green.

**Figure 2-figure supplement 3.**
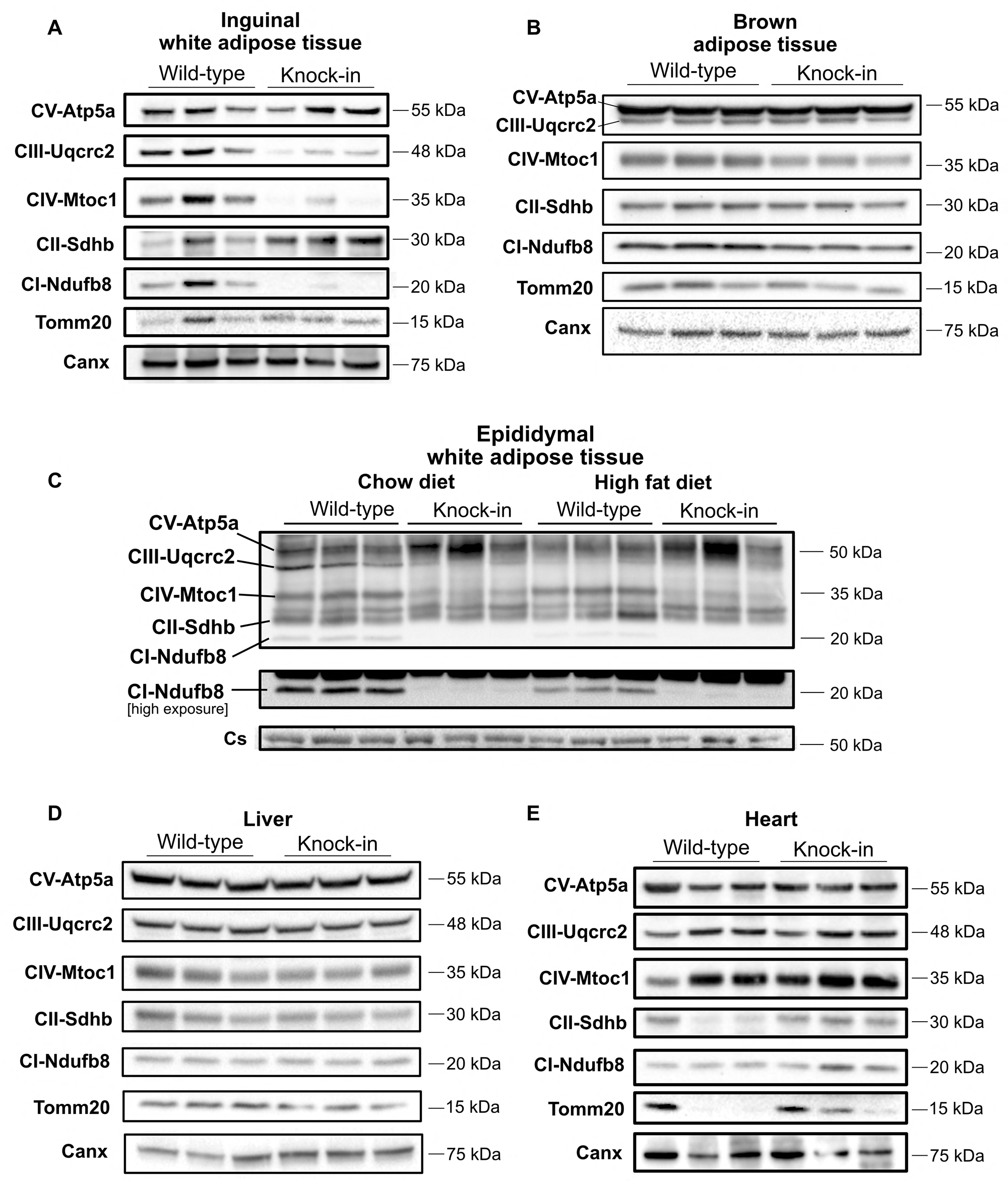
Altered expression of Oxphos protein subunits in adipose tissue. Western blots showing expression of Oxphos protein subunit expression in mice fed chow for 6 months. Expression is shown from inguinal white adipose tissue (A), brown adipose tissue (B), epididymal white adipose tissue (C), liver (D), and heart (E). Each lane contains samples from a separate animal and blots are representative of at least three replicates. *Canx* (calnexin) is used as total cellular control and *Tomm20* (or citrate synthetase (*Cs*) is used as mitochondrial mass loading control. **Figure 2-figure supplement 3-source data.** Raw and annotated Western blots for Oxphos subunits, Tomm20, calnexin, and citrate synthetase for all tissues.

**Figure 2-figure supplement 4.**
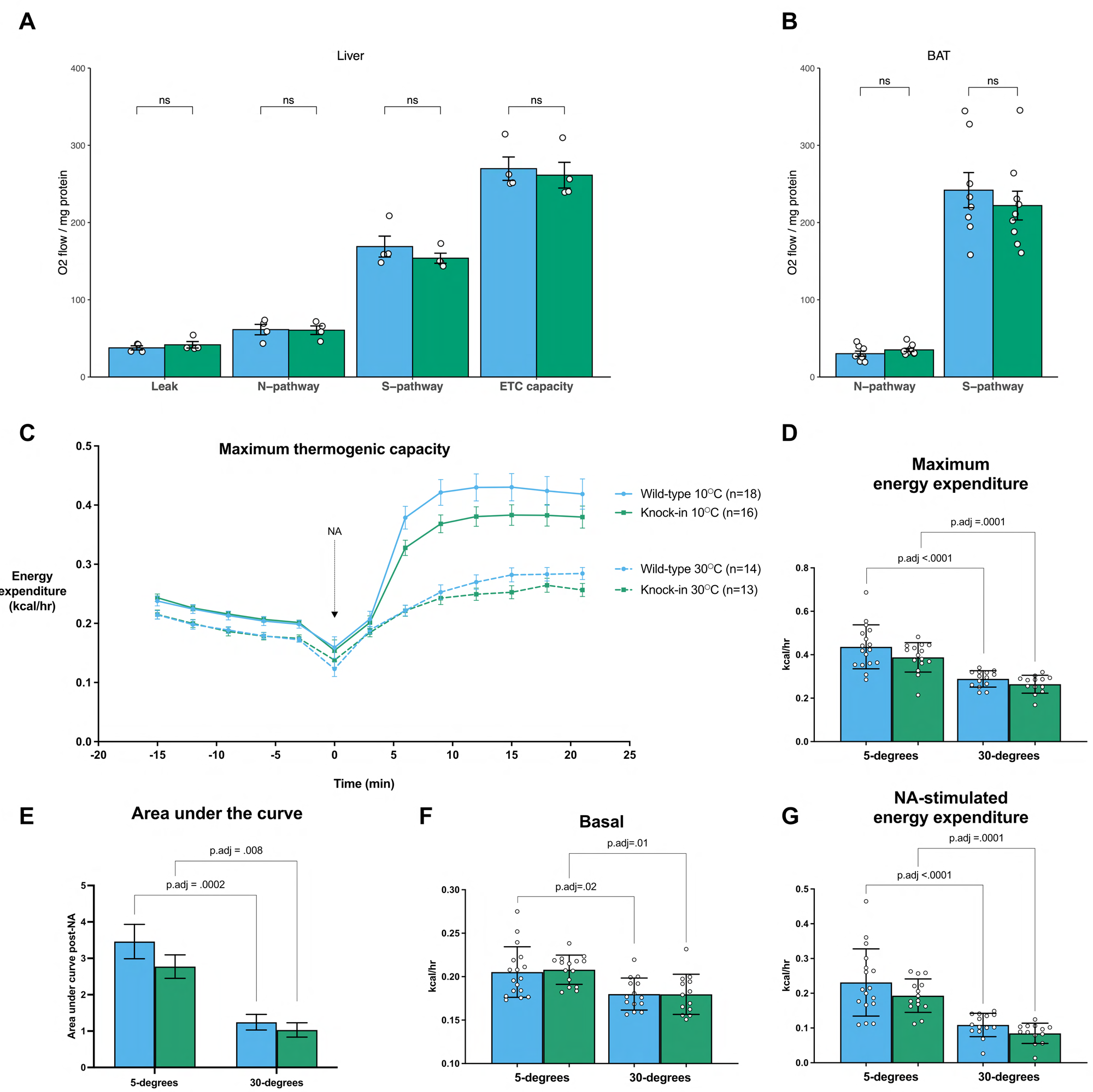
Mfn2^R707W^ does not impair brown adipose tissue thermogenic capacity. *Ex vivo* mitochondrial respirometry in liver (A, n=4) and BAT (B, n=9). Each data point represents data from a separate animal. Eight-week-old mice were exposed to cold (10°C, n=16-18) or thermoneutrality (30°C, n=13-14) for 4 weeks then maximum thermogenic capacity was tested using noradrenaline (NA) stimulation under anaesthesia. (C) Trend in energy expenditure (kcal/hr) before and after noradrenaline stimulation for cold (solid line) and thermoneutral (dashed line) housed animals. (D) Quantification of maximum (peak) energy expenditure. P-value represents the difference between WT groups comparing 10°C and 30°C. (E) Area under curve analysis using the minimum value at the point of noradrenaline (NA) injection at the base of the curve. (F) Basal energy expenditure prior to NA injection. (G) Difference between peak and baseline energy expenditure under both conditions. WT in blue, homozygous Mfn2^R707W^ in green. Each data point represents results from an individual animal.

**Figure 3-Figure supplement 1.**
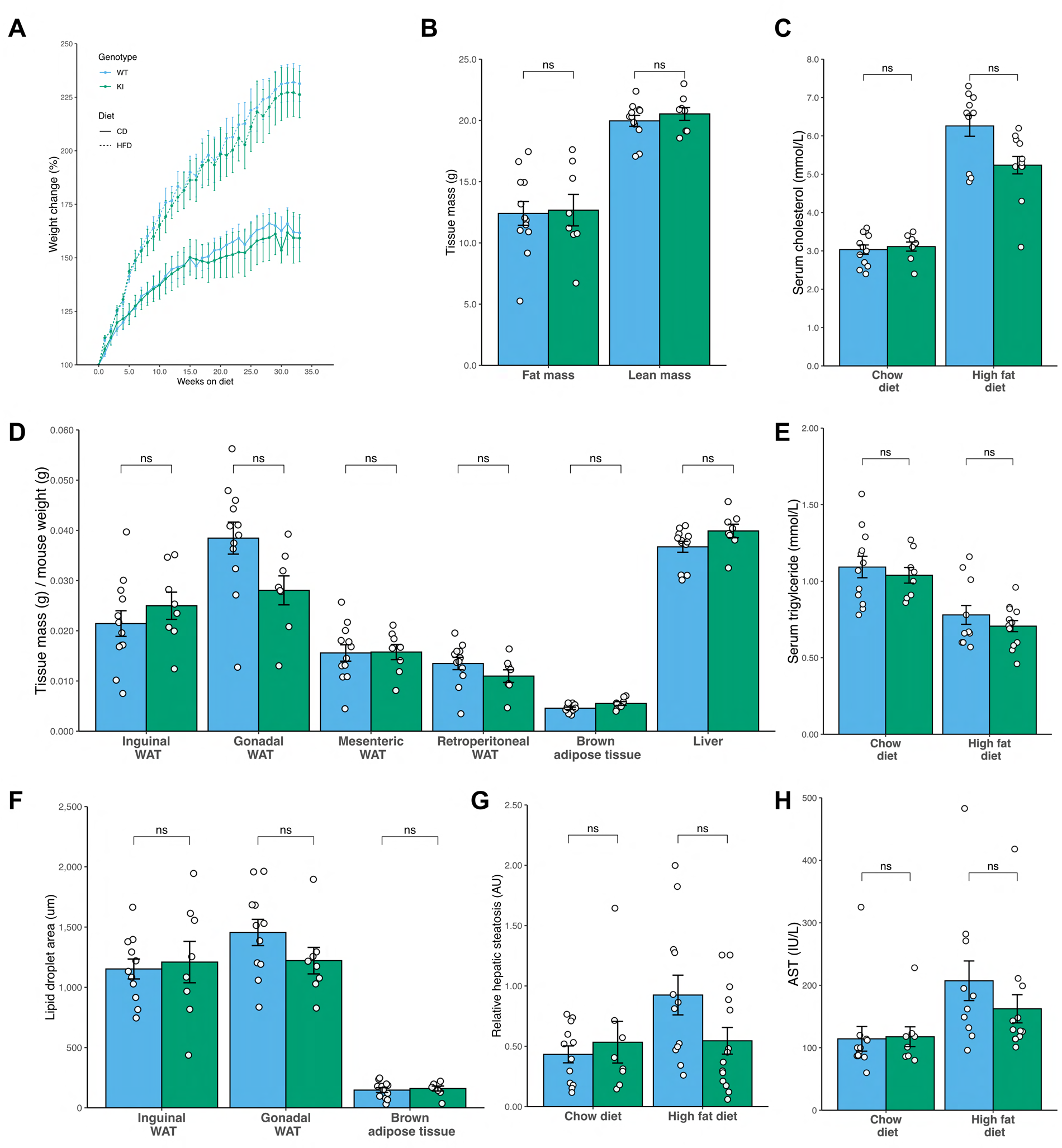
No evidence of altered fat mass or glucose homeostasis in Mfn2^R707W^ mice on chow or high fat diet for 6 months. Seven-week-old mice were fed chow diet (n=8-12) or 45% kcal HFD (n=13-14) for 6 months. (A) Relative (%) change body mass for mice fed chow diet (solid line) and HFD (dashed line) over 6 months. (B) Time-domain nuclear magnetic resonance (TD-NMR) measurement of fat and lean mass in mice fed a chow diet 6 months. (C) Analysis of mouse serum biochemistry after 6 months of diet and a 6 hr fast for total cholesterol. (D) Weights of tissues, including four white adipose tissue (WAT) depots, from mice fed chow diet for 6 months. (E) Serum triglycerides. (F) Quantification of lipid droplet area from histological specimens of adipose tissue from mice fed chow diet for 6 months. (G) Quantification of relative hepatic steatosis from histological images of liver. (H) Serum aspartate aminotransferase. ns, p>.05 on un-paired T-test, adjusted for multiple comparisons. WT in blue, homozygous Mfn2^R707W^ in green. Each data point represents results from an individual animal.

**Figure 4-figure supplement 1.**
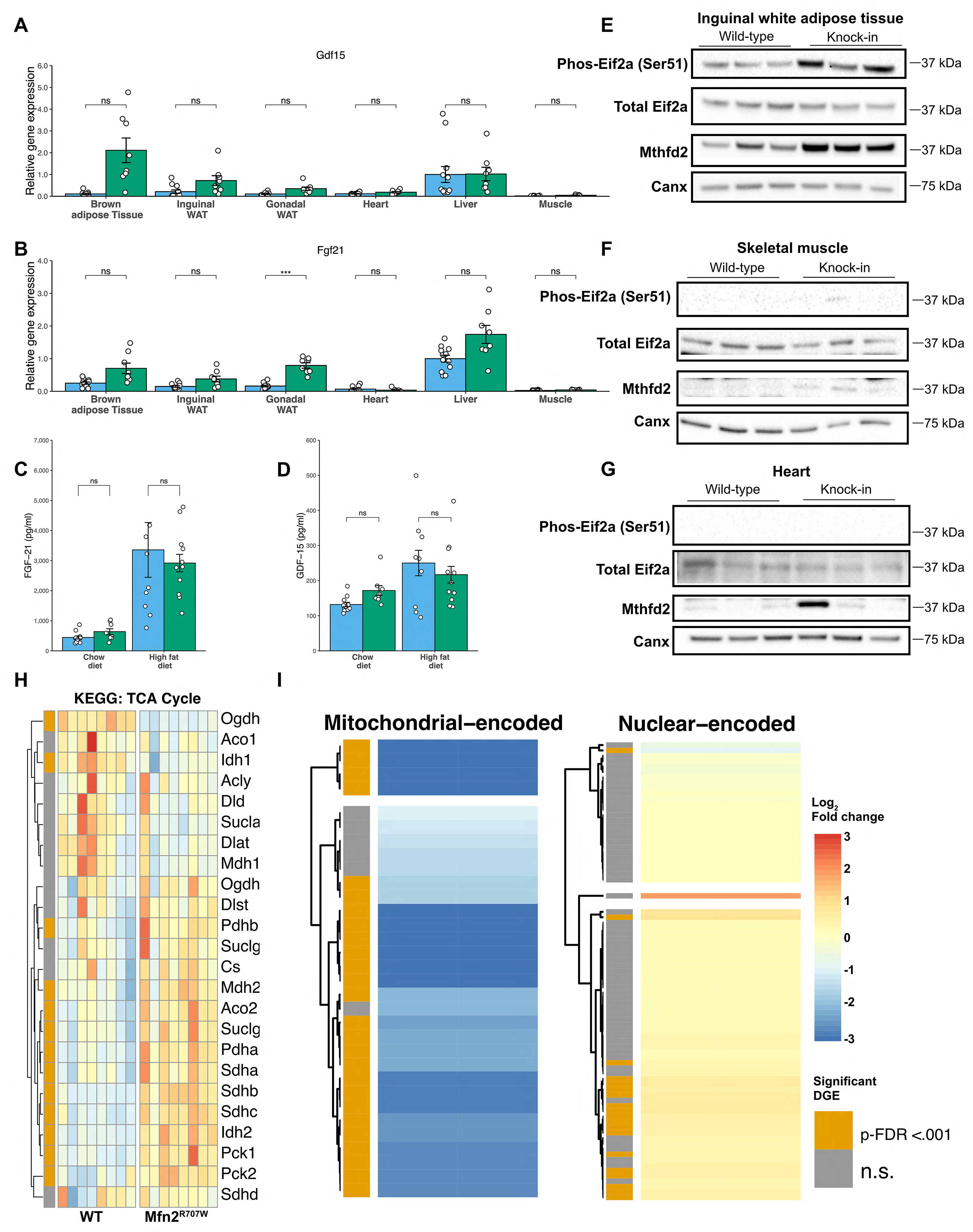
Mfn2^R707W^ causes adipose tissue-specific induction of the integrated stress response with perturbation of mitochondrial gene expression. qPCR of genes involved in the integrated stress response (ISR) for six tissues from animals fed a chow diet for 6 months: *Gdf15* (A) and *Fgf21* (B). Each data point represents data from a separate animal. Target gene CT values were normalised against three housekeeping genes (*36b4, B2m,* and *Hprt*) and expressed relative to WT liver for each gene. p-values are FDR-adjusted for multiple tests. Serum GDF-15 (C) and FGF-21 (D) after 6 months of chow or HFD. Western blots from inguinal white adipose tissue (E), skeletal muscle (F), and heart (G) illustrating Ser51-phosphorylation of eIF2a and expression of Mthfd2 with calnexin (*Canx*) as loading control. Western blots are representative of at least three biological replicates. (H) Heatmap of all genes from the citrate acid (tricarboxylic acid (TCA)) cycle KEGG pathway. Colour illustrates the Log_2_ fold change normalised per gene. Those annotated with orange in the leftmost column had significant differential gene expression (DGE). (I) Heatmaps comparing mRNA expression of mitochondrial- and nuclear-encoded mitochondrial genes from inguinal WAT from HFD-fed animals. *** p-FDR <.001. **Figure 4-figure supplement 1-source data.** Raw and annotated Western blots for Phospho-/Total-Eif2a, Mthfd2, and loading calnexin from panels E, F, and G.

**Figure 4-figure supplement 2.**
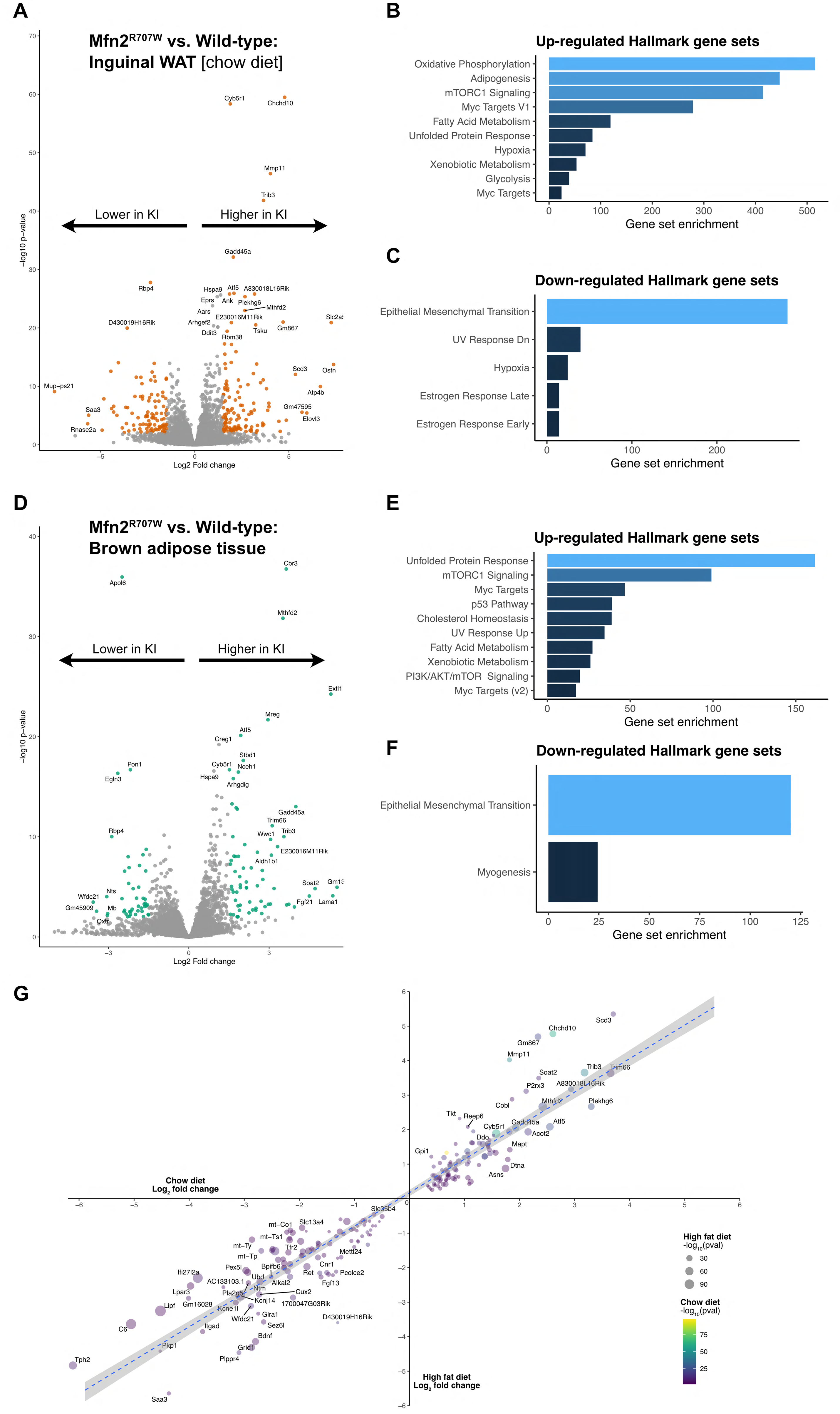
Transcriptional evidence of upregulation of the unfolded protein response and mTorc1 pathways in adipose tissue. (A) Volcano plot from bulk RNA sequencing (n=7 WT and n=6 KI) of inguinal white adipose tissue from mice on chow, where significantly differentially expressed genes (Log_2_ fold change >1.5 and p-FDR <.001) are highlighted in green. Pathway analysis using significantly differentially expressed genes for upregulated (B) and downregulated (C) Hallmark gene sets. (D) Volcano plot from bulk RNA sequencing (n=6 per genotype) of brown adipose tissue from mice on HFD, where significantly differentially expressed genes (Log_2_ fold change >1.5 and p-FDR <.001) are highlighted in green. Pathway analysis using significantly differentially expressed genes for upregulated (E) and downregulated (F) Hallmark gene sets. All illustrated gene sets are enriched with p-FDR <.05. (G) Scatter plot of the top 100 up-/down-regulated genes from bulk RNA sequencing from inguinal white adipose tissue. For each gene, the fold change observed under HFD is plotted on the x-axis and fold change under chow diet is shown on the y-axis. P-values for each dietary condition are shown using size (HFD) and colour (chow diet).

**Figure 5-figure supplement 1.**
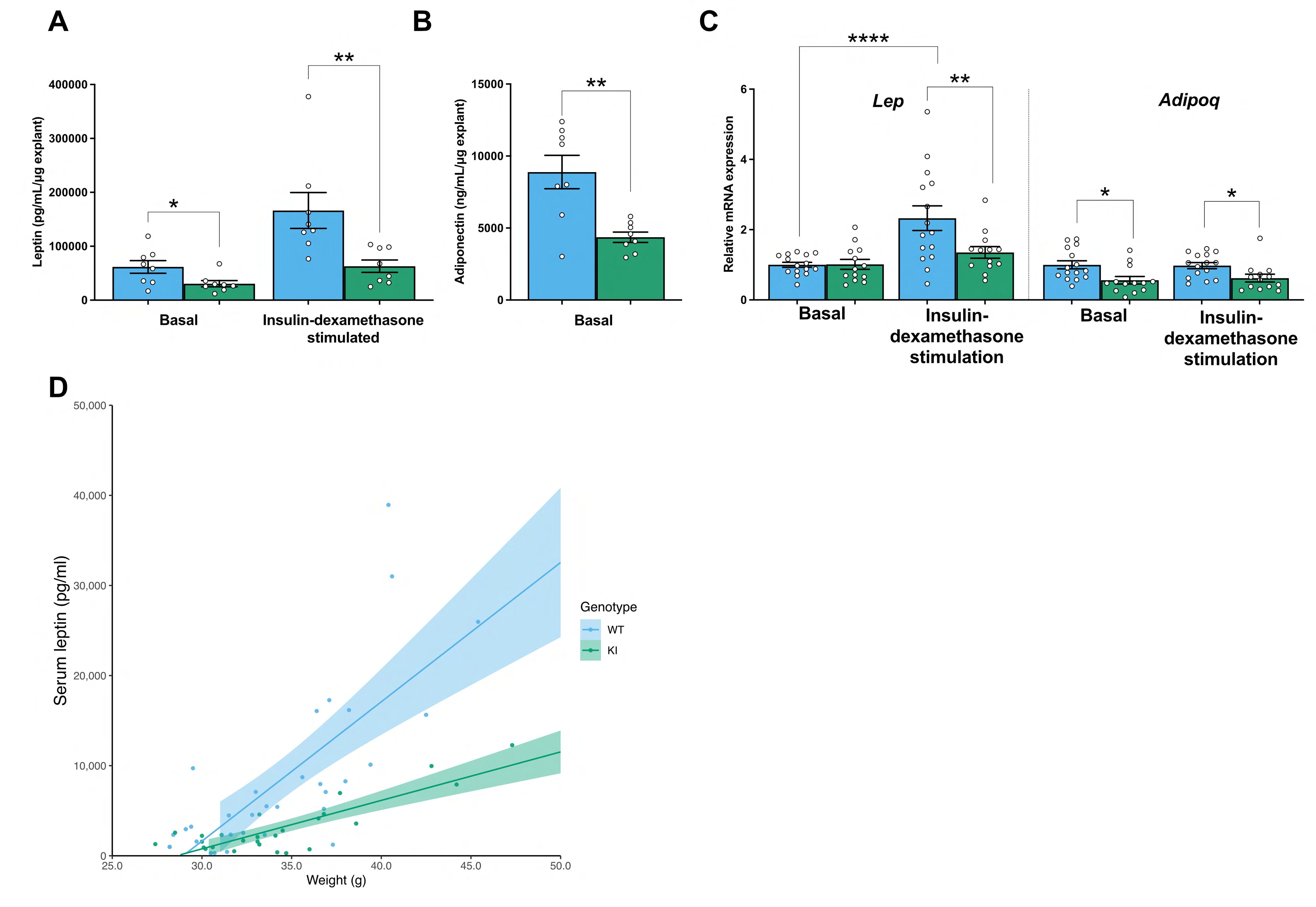
Mfn2^R707W^ decreases adipose leptin and adiponectin secretion from multiple adipose depots and in different dietary conditions. (A) Leptin secretion from explants of epididymal white adipose tissue in basal conditions and after insulin-dexamethasone stimulation. (B) Adiponectin secretion from epididymal WAT explants. (C) qPCR of *Lep* and *Adipoq* from epididymal explants from animals fed a HFD, where each data point represents an explant from a separate animal (n=11-13). (D) Relationship between leptin and body weight for animals on chow. Each data point represents one measurement of leptin, with multiple measurements per animal. Shaded area represents the 95% confidence interval. Data from n=8-12 animals. Asterisks indicate p-values are FDR-adjusted for multiple tests (* p-FDR <.05, ** p-FDR <.01, *** p-FDR <.001).

## Tables

**Supplementary Table 1:**
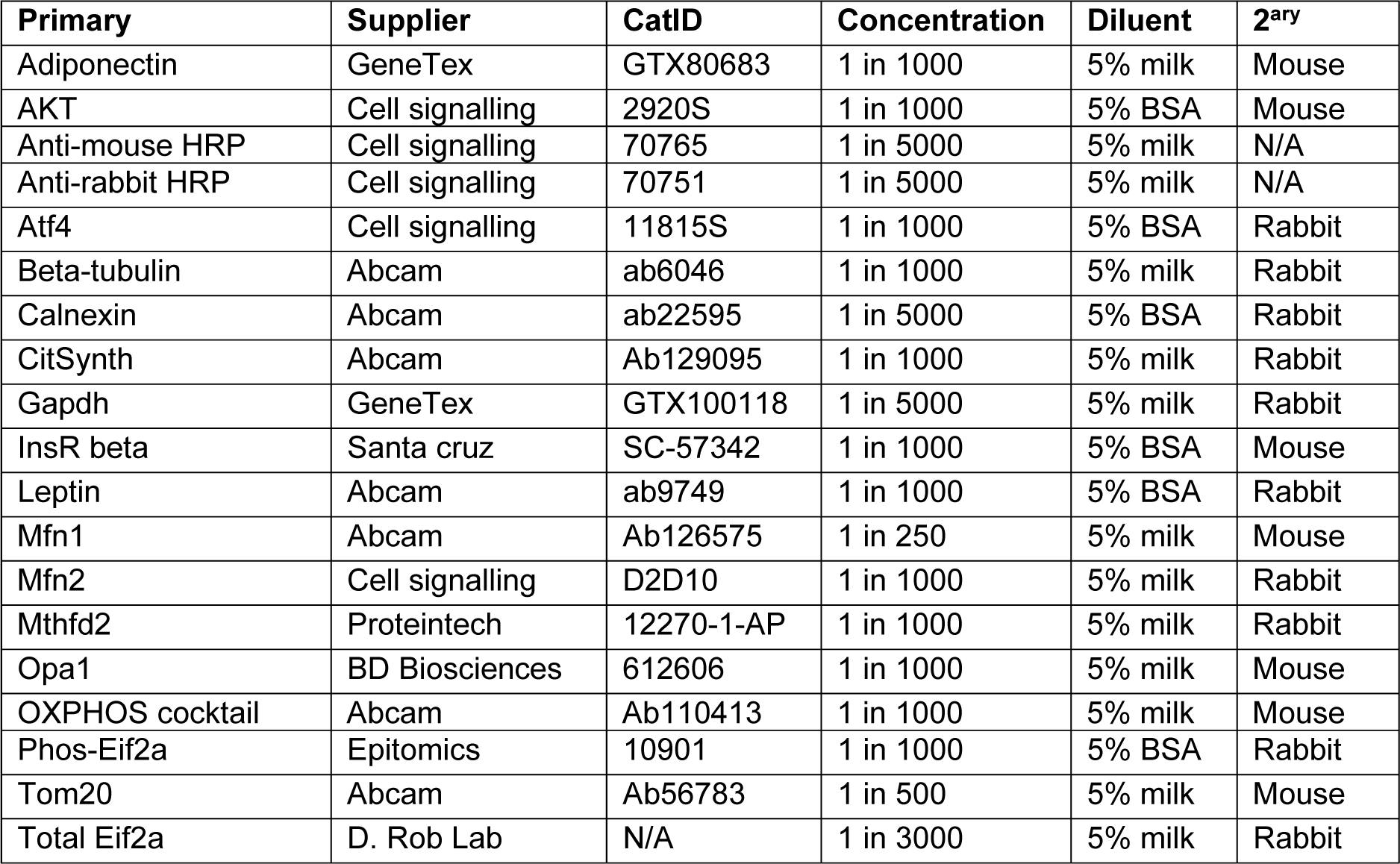
Antibotides used in this study.

**Supplementary Table 2:**
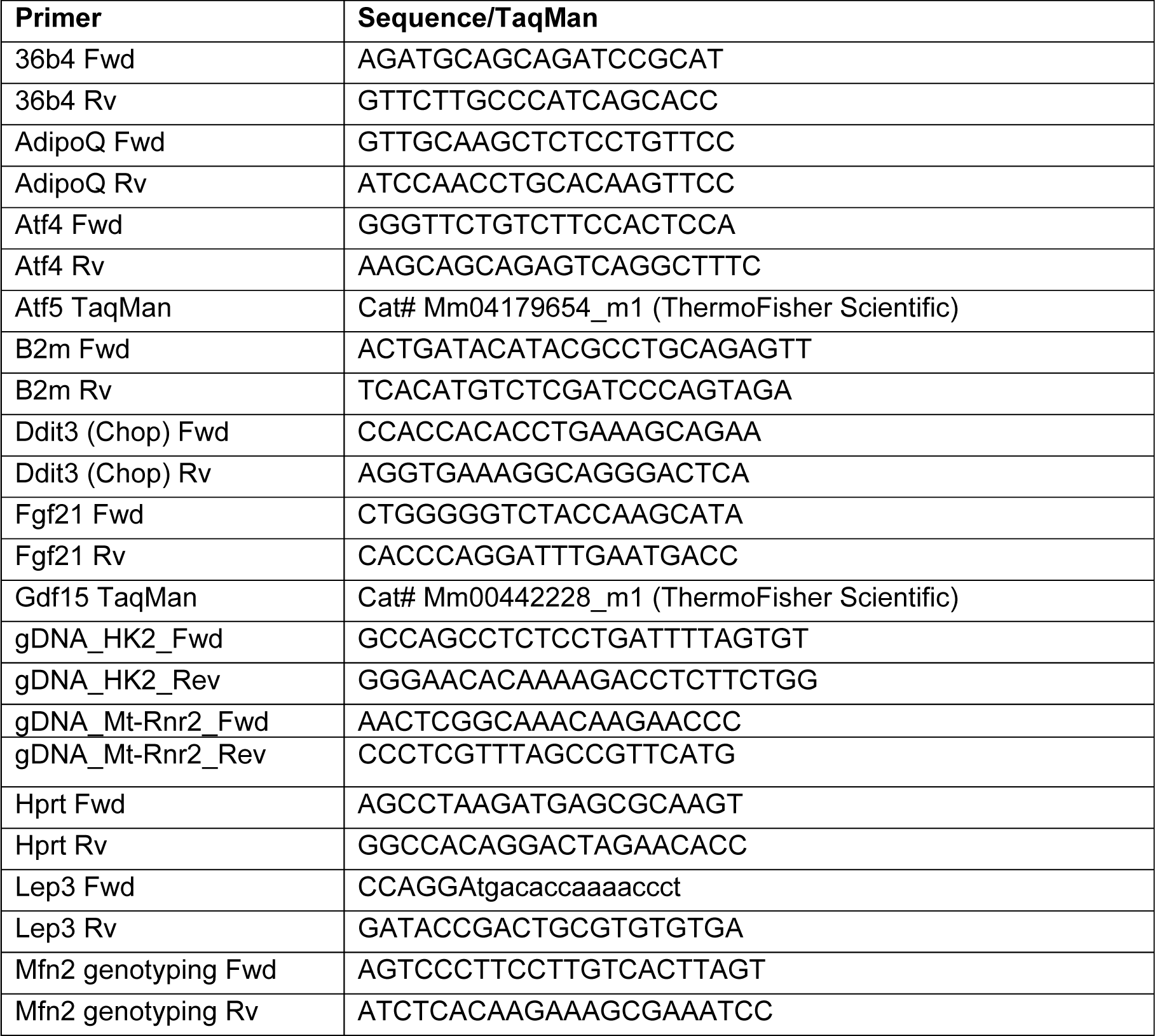
Primer sequences used in this study. Fwd, forward primer; Rv, reverse primer.

